# Oligomerization enables the selective targeting of intrinsically disordered regions by small molecules

**DOI:** 10.1101/2025.03.21.644603

**Authors:** Stasė Bielskutė, Borja Mateos, Muhammad Awawdy, Carla Garcia-Cabau, Henri Niskanen, Carolina Sánchez-Zarzalejo, Lorenzo Bracaglia, Roberta Pierattelli, Isabella C. Felli, Marta Frigolé-Vivas, Jesús García, Antoni Riera, Denes Hnisz, Xavier Salvatella

## Abstract

Intrinsically disordered regions (IDRs) are challenging drug targets because they lack stable interaction sites for drug-like molecules. We studied the first small molecule targeting an IDR to be evaluated in a clinical trial and found that it interacts selectively with an oligomeric form of the protein, more structured than the monomeric state, that is stabilized by interactions involving aromatic residues in regions with helical propensity. We also found that this compound alters the network of interactions defining the conformational ensemble of its target, thus affecting its condensation properties, linked to its function as a transcription factor. These findings provide a framework for developing strategies to target intrinsically disordered regions with small molecules.

## Introduction

The intrinsically disordered regions (IDRs) of proteins are highly attractive targets for therapeutic intervention due to their diverse biological activities and functional versatility^1^. However, these regions are often considered undruggable because they lack stable secondary and tertiary structure, which are required to form well-defined interaction sites for drug-like small molecules^2,3^. Despite this challenge, a handful of compounds capable of interacting with IDRs have been identified but the extent to which these interactions are sufficiently selective to enable drug development remains poorly understood^4–9^.

To address this important question, we investigated the selectivity of the interaction between the first drug-like small molecule targeting an IDR to reach clinical trials, EPI-001, and its target, the activation domain (AD) of the androgen receptor (AR)^10,11^. We found that EPI-001, initially identified in a phenotypic screen, interacts selectively with an oligomeric form of the AR AD, which has a higher degree of structuration than the monomer and is stabilized by interactions between aromatic residues in partially helical regions of sequence.

Our findings also revealed that the interaction of EPI-001 with the AR AD rewires the conformational ensemble of the target, alters the physical properties of the AR AD phase-separated state and reduces its ability to recruit RNA Polymerase II (RNAPII). These results indicate that IDRs can be selectively targeted with drug-like small molecules if they transiently form druggable conformational states. In summary, our results show that it is possible to discover and develop small molecule inhibitors of this class of highly challenging drug targets.

## Results

### EPI-001 selectively interacts with oligomeric AR AD

To assess the selectivity of EPI-001 towards the AR, we examined its interaction with the AR AD and the ADs of four other nuclear receptors (NRs): the mineralocorticoid (MR), progesterone (PR), glucocorticoid (GR), and estrogen (ERɑ) receptors (Fig. 1a)^12,13^. NRs are transcription factors with conserved domain structures that serve as drug targets across various disease areas^14,15^. Their IDRs were chosen as a stringent selectivity panel due to their comparable functional roles and sequence properties. While sequence alignment alone does not immediately reveal these similarities (Fig. 1b), they become clear through parameters such as amino acid composition (Fig. 1c) and sequence patterning (Fig. 1d and Extended Data Fig. 1a,b), that differ from those of IDRs not belonging to the panel such as hnRNPA1 and FUS (Extended Data Fig. 1c)^16–18^. Solution nuclear magnetic resonance (NMR) spectroscopy revealed that the NR ADs displayed low chemical shift dispersion and broad dynamic ranges, consistent with the behavior of IDRs enriched in transient long-range interactions (Extended Data Fig. 1d)^19–22^. Unlike the AR AD, where EPI-001 addition caused chemical shift perturbations (CSPs) (Fig. 1e, Extended Data Fig. 1e), those detected for the other members of the selectivity panel were much weaker (Fig. 1e)^9,23^. Notably, a large number of resonances in the spectrum of the MR AD were broadened beyond detection. Therefore, to rule out a potential interaction of EPI-001 with the NMR undetectable regions of MR AD, we added MR AD to an AR AD with EPI-001 solution and observed no competition (Extended Data Fig. 1f,g). Although CSPs are not absolute measures of intermolecular interaction strength, our findings—together with the CSPs induced by NR ADs on EPI-001 (Fig. 1f, Extended Data Fig. 1h)—indicate that EPI-001 is a selective ligand of the AR AD.

**Fig. 1.**
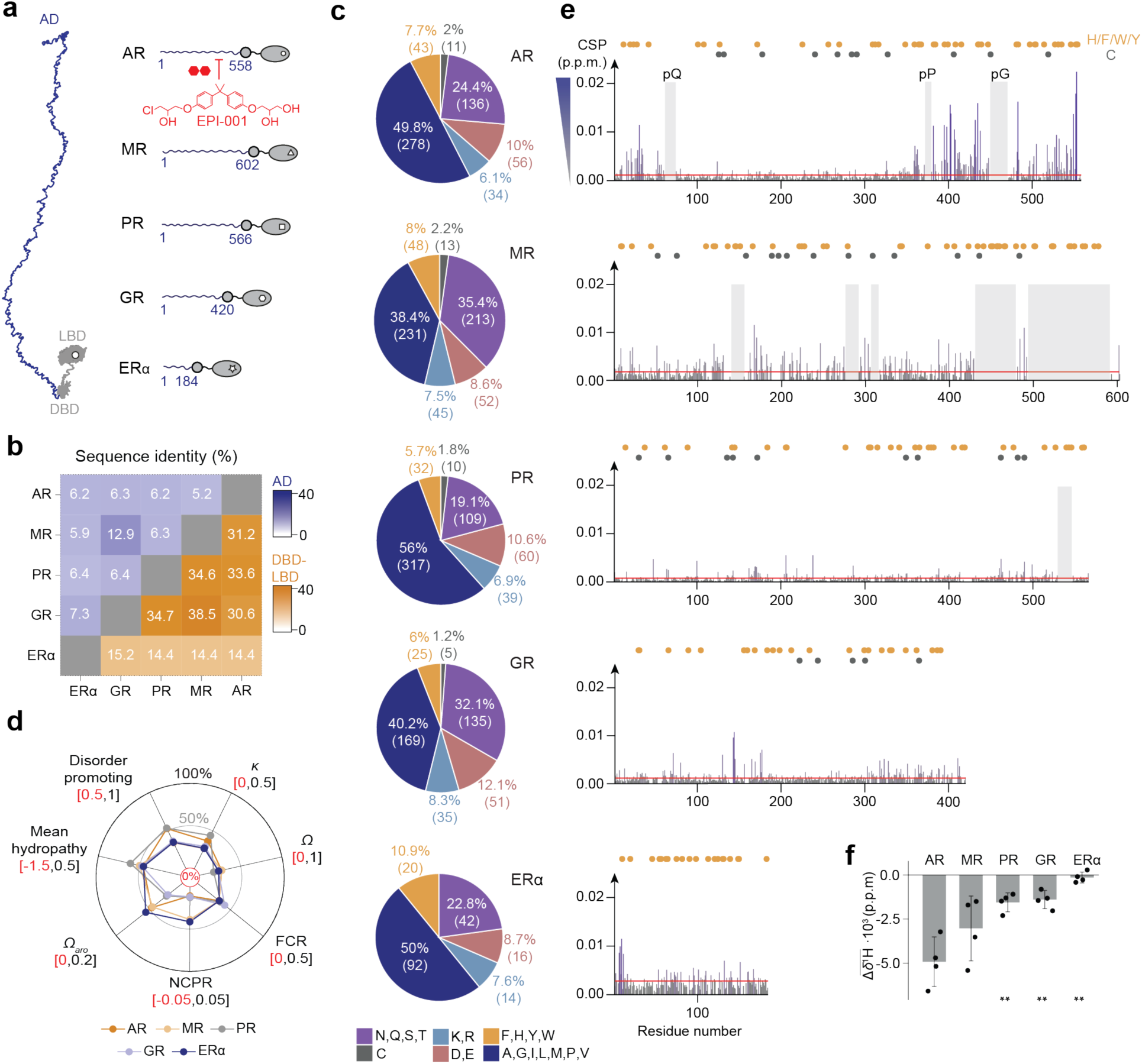
EPI-001 interacts selectively with the activation domain of AR. (**a**) The selectivity panel is composed of nuclear receptors activated by steroid hormones, possessing a long variable AD followed by conserved globular domains (DBD and LBD). (**b**) Sequence identity fraction of the ADs and DBD-LBD in blue and orange, respectively. (**c**) Aminoacid composition fractions for five different residue types (polar - purple; cysteine - grey; positive charge - light blue; negative charge - pink; aromatic - orange; hydrophobic - blue). (**d**) Overall sequence properties for the different activation domains. *κ* and *Ω* values describe the segregation of charged and proline residues, respectively; FCR, fraction of charged residues; NCPR, net charge per residue; and *Ω_aro_* represents the aromatic clustering. (**e**) CSPs extracted from 2D ^1^H-^15^N NMR correlation spectra of the corresponding AD induced in the presence of 10 molar equivalents of EPI-001. Orange and grey circles indicate the position of aromatic and cysteine residues, respectively. The red line represents the significant threshold calculated as the average plus five standard deviations of the first quartile of CSPs. Grey shaded boxes represent the unassignable regions in the NMR spectra because of the broadening beyond detection of some protein resonances or due to signal overlapping on some homorepeat regions. (**f**) Average ^1^H shifts from distinctive EPI-001 signals in the presence of 0.1 molar equivalents of the corresponding AD. The error bar denotes the standard deviation (*n* = 4).

EPI-001 perturbs resonances of the AR AD^9,23^ that decrease in intensity upon AR AD oligomerization^23,24^. A comparison of these two datasets revealed a correlation (R² = 0.7, Fig. 2a), suggesting that the CSPs caused by EPI-001 could be, at least partially, indirect and the result of an increased population of AR AD oligomers. We therefore used dynamic light scattering (DLS) to analyze the oligomerization propensity of the NR ADs. We obtained that the AR, ERɑ, GR, and PR ADs had similar oligomerization propensities, all significantly lower than that of the MR AD (Extended Data Fig. 2a-c), explaining the line broadening observed in the MR AD spectra (Fig. 1e)^23^. The population of AR AD monomers decreases monotonically upon increasing the concentration of EPI-001 (Fig. 2b) but the oligomerization equilibrium of the MR AD, the member of the selectivity panel with the highest oligomerization propensity, is not affected by EPI-001. Those of the other members of the selectivity panel are affected but to a lesser extent than that of the AR AD (Extended Data Fig. 2d). These results indicate that EPI-001 selectively stabilizes the oligomers formed by the AR AD.

**Fig. 2.**
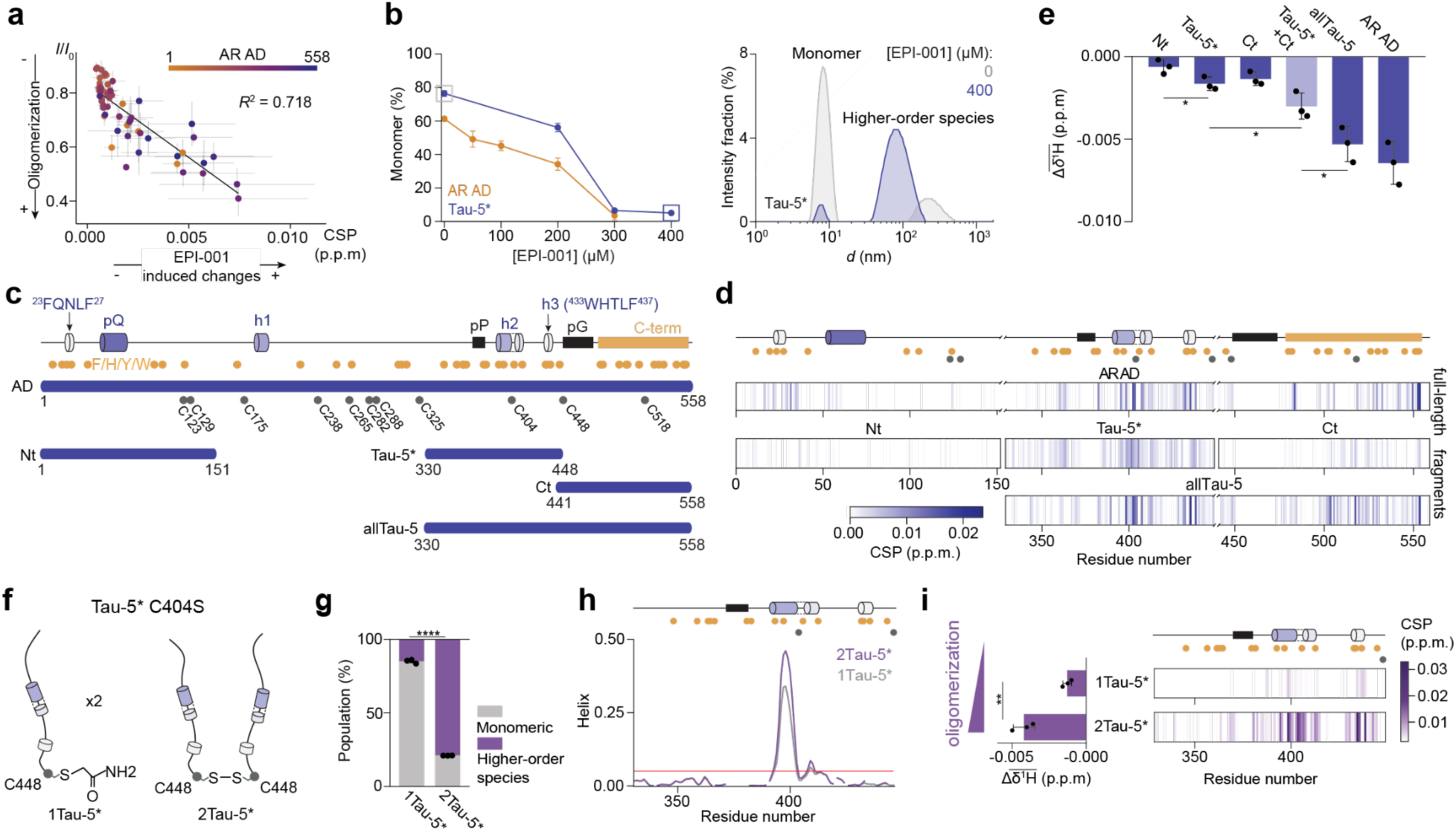
EPI-001 interacts with oligomeric AR AD. (**a**) Correlation between the regions involved in AR AD oligomerization^23^ and the perturbed ones upon the addition of EPI-001. (**b**) (*Left*) Percentage of monomer in 35 µM AR AD and 400 µM Tau-5* samples at varying concentrations of EPI-001. The remaining percentage represents high-order species measured by DLS. (*Right*) DLS intensity measurement of Tau-5* in the absence (gray) and presence (blue) of 400 µM EPI-001. (**c**) Schematic illustrating the AR AD fragments designed. Helical motifs are represented as cylinders (Extended Data Fig. 2f) and proline and glycine homorepeats as black squares. Orange and grey circles indicate aromatic and cysteine residues, respectively. (**d**) CSPs are depicted in barcode plots extracted from 2D ^1^H-^15^N NMR correlation spectra of 25 µM AR AD and its fragments in the absence and presence of 250 µM of EPI-001. (**e**) Average of the ^1^H shifts induced on three distinct EPI-001 signals in the presence of 0.1 molar equivalent of AR AD and its fragments. The Tau-5*+Ct lighter blue bar represents the sum of the independent Tau-5* and Ct experiments. (**f**) 2Tau-5* is formed by a disulfide bond at C448, while in 1Tau-5*, C448 is blocked with iodoacetamide to prevent disulfide bond formation. (**g**) Percentage of monomeric and high-order species in 1Tau-5* and 2Tau-5* samples measured by DLS at a concentration of 2.44 mg/mL. (**h**) NMR-derived helical content of 1Tau-5* and 2Tau-5* at 0.31 mg/mL. The red line indicates the uncertainty of the *δ*2D algorithm (5%)^51^. (**i**) *(Left)* Average shifts of 250 µM EPI-001 ^1^H signals induced by 0.31 mg/mL of 1Tau-5* or 2Tau-5*. *(Right)* Per-residue CSPs in 2D ^1^H-^15^N NMR correlation spectra of 0.31 mg/mL 1Tau-5* or 2Tau-5* after adding 250 µM EPI-001. In both cases, the ratio corresponds to 1:10 of EPI-001 interaction sites on the protein to EPI-001. (**e**,**g**,**i**) Error bars indicate standard deviations (*n* = 3).

We next carried out experiments to identify the site of interaction of EPI-001 on the AR AD. Since target CSPs appear to be a convolution of direct chemical shifts—caused by contacts between target and ligand—and indirect ones—caused by changes in the structure or degree of oligomerization, we studied the CSPs induced by EPI-001 on AR AD shorter variants Nt (residues 1-151)^25^, Tau-5* (residues 330-448)^9^, and Ct (residues 441-558)^23^ (Fig. 2c and Extended Data Fig. 2e,f), which encompass the regions where CSPs are observed (Fig. 1e), as well as allTau-5 (residues 330-558), which spans Tau-5* and Ct (Fig. 2c). We found that EPI-001 caused CSPs only in Tau-5* and allTau-5 (Fig. 2d,e and Extended Data Fig. 2g,h), indicating that Tau-5* contains the site of interaction of this small molecule and, consequently, that the CSPs observed in Nt and Ct are mainly indirect.

That EPI-001 stabilizes an oligomeric form of the AR AD suggests that it interacts with it. To investigate this hypothesis we generated a chimeric form of Tau-5* C404S by forming a disulfide bond between the side chains of the C-terminal Cys residues (C448) of two different Tau-5* C404S molecules, called 2Tau-5*, and compared its properties to those of 1Tau-5*, which we obtained by reacting with iodoacetamide the Cys 448 side chain of Tau-5*C404S (Fig. 2f). First we used DLS to compare the oligomerization propensity of 1Tau-5* and 2Tau-5*. We found that, at the same mass concentration, 2Tau-5* had a higher propensity to form high order oligomers than 1Tau-5* (Fig. 2g and Extended Data Fig. 2i), presumably because of the increase in multivalency^26^. Next we analyzed the chemical shift differences between 1Tau-5* and 2Tau-5* (Extended Data Fig. 2j,k), which were equivalent to those induced by increasing the protein concentration of Tau-5*^24^, in agreement with the DLS results. In addition, the ^13^Cα and ^13^C’ chemical shifts indicated that oligomerization increases the helicity of helix h2 (Fig. 2h).

Our results indicate that introducing a covalent bond between C-terminal Cys residues increases the oligomerization propensity of Tau-5* C404S while preserving the site of interaction of EPI- 001, providing us with an opportunity to test that EPI-001 has a higher affinity for the oligomeric than for the monomeric form of the target. In agreement with this hypothesis, larger CSPs were observed in 2Tau-5* than 1Tau-5*in the presence of EPI-001 (Fig. 2i and Extended Data Fig. 2g,k,l). Finally we used microscale thermophoresis (MST) to verify that EPI-001 interacts with 2Tau-5* with an orthogonal biophysical technique (Extended Data Fig. 2m). We concluded that EPI-001 interacts with an oligomeric form of AR AD that is more structured than the monomer.

### EPI-001 rewires the network of interactions of the AR AD

To investigate the network of homo and heterotypic interactions defining the conformational ensemble of the AR AD we measured the CSPs induced by constructs Nt, Tau-5* and Ct on the resonances of the same constructs in equimolar solutions (Extended Data 3a). To summarise the information contained in these 9 experiments we computed for pairs of residues a contact parameter, *σ_ij_*^apo^, as the product of the CSPs measured for residues *i* and *j* in the two reciprocal experiments, obtaining the CSP matrix shown in Fig. 3a; for consistency, *σ_ii_*^apo^ was computed as the square of the CSP observed for residue *i*. We found that pairs of residues in motifs ^23^FQNLF^27^ and ^433^WHTLF^437^ have overall high *σ*^apo^ values, suggestive of a direct interaction between these motifs, and a similar result was obtained for the former motif and regions rich in aromatic residues in Ct and Tau-5*, although with lower values (Fig. 3a and Extended Data Fig. 3a). To validate that high values of *σ*^apo^ reflect the network of interactions we used paramagnetic relaxation enhancement (PRE) NMR experiments. Specifically, we used these to measure intra- and intermolecular interactions involving C404 in the full-length AR AD, as this native Cys residue is found in the site of interaction of EPI-001 (Extended Data Fig. 3b). We found the regions identified by the CSP matrix are involved in both intra- and intermolecular interactions, confirming that the CSP matrix reports on the interactions of full-length AR AD.

**Fig. 3.**
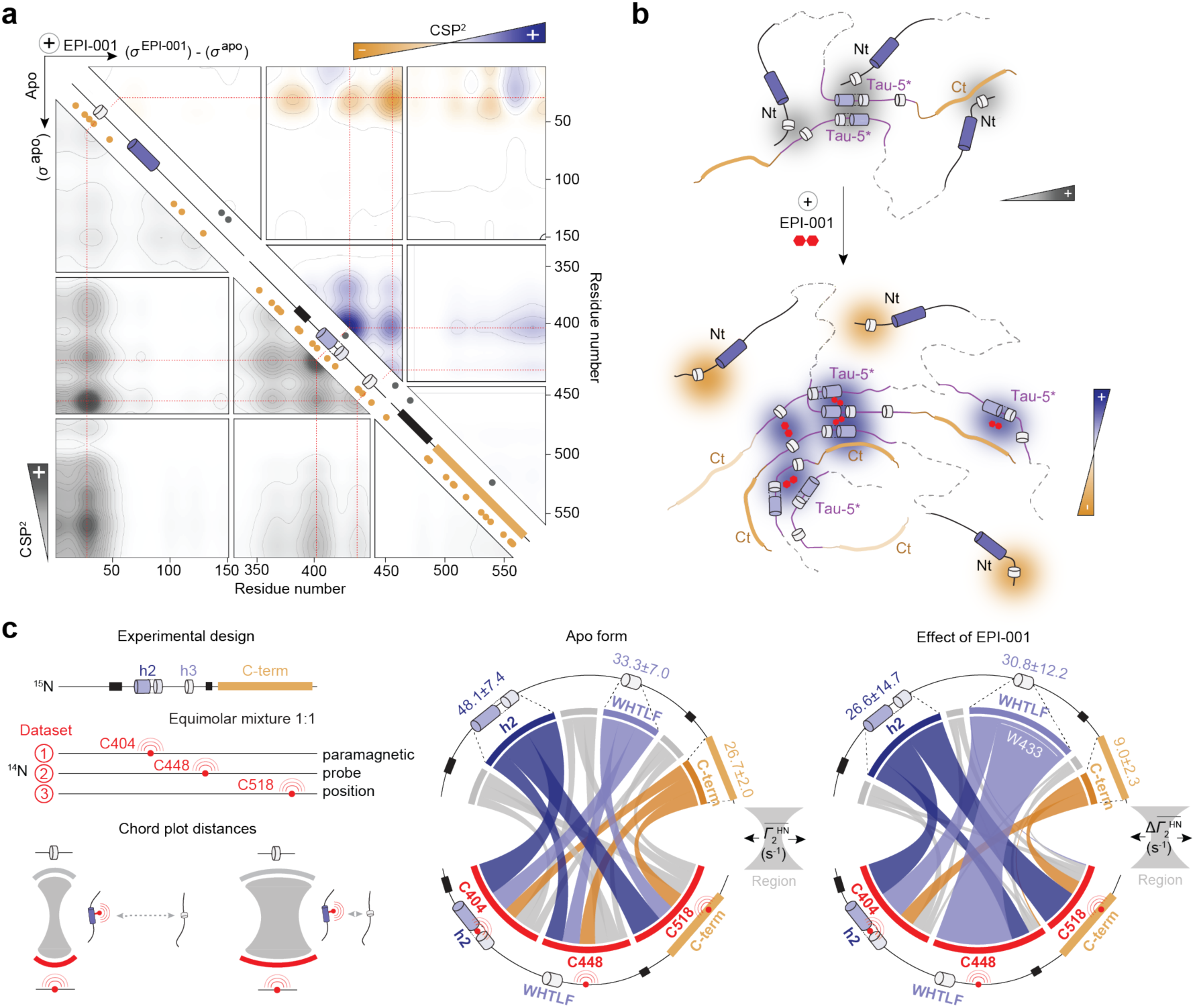
EPI-001 rewires the network of interactions within AR AD and strengthens homotypic interactions between residues in Tau-5*. (**a**) CSP matrix between AR fragments in equimolar mixtures and in the absence of EPI-001 is shown in the lower matrix. Differences between AR fragments in the presence and absence of 1 molar equivalent EPI-001 are shown in the upper matrix. Red dashed lines represent helical motifs enriched in aromatic residues. Orange and grey dots indicate the position of aromatic or cysteine residues, respectively. (**b**) Schematic representing interactions between AR AD regions and the changes induced by EPI-001. (**c**) Intermolecular interactions between allTau-5 as monitored by PRE experiments. *(Top)* Experimental design *(left)* and interpretation *of Γ*_2_^HN^ rates (*right*). (*Bottom*) Chord diagram^52^ representing the strength of intermolecular contact. The chord width indicates the average *Γ*_2_^HN^ of contact between the corresponding regions in the absence of EPI-001 (*left*) or the average Δ*Γ*_2_^HN^ induced by adding 1 molar equivalent of EPI-001 (*right*). The average *Γ*_2_^HN^ for certain regions are shown above the plots. W433 contributes significantly to the average. (Extended Data Fig. 3h).

The CSP matrix also reveals the relative self-association propensity of the fragments: Nt and Ct appear to have lower *σ*^apo^ values, indicating a lower propensity to oligomerize than Tau-5*^24^ (Fig. 3a and Extended Data Fig. 3a). Given that this fragment and Ct have similar aromatic character (Extended Data Fig. 2e), this result suggests that the conformational properties of the fragment such as its propensity to form secondary structures (Extended Data Fig. 2f), influence its propensity to oligomerize. In line with this, the motifs with some of the highest *σ*^apo^ values have high helical propensity or fold into ɑ-helices upon binding to globular partners^27,28^ (Extended Data Fig. 2f). To confirm this we re-measured the CSP matrix after introducing three helix-breaking substitutions (L26P in Nt, A398P and L436P in Tau-5*) and observed lower and less well-defined *σ*^apo^ values, reflecting weaker, less specific interactions, and in line with the notion that aromatic residues in helices are particularly prone to be involved in interactions (Extended Data Fig. 3c,d).

We analysed how this drug-like small molecule influences the CSP matrix to characterize the population shifts caused by EPI-001 on the AR AD . We found that EPI-001 appears to strengthen (Δ*σ*>0, blue) the homotypic interactions between the region of sequence centred around residue 400, partially helical and rich in aromatic residues, and instead weakens (Δ*σ*<0, orange) heterotypic interactions between the motif ^23^FQNLF^27^ and the rest of the sequence (Fig. 3a, b and Extended Data Fig. 3a). Based on this result we hypothesized that EPI-001 competes with the ^23^FQNLF^27^ motif for interaction with the helical motifs h2 and h3 in Tau-5*. To test this we used variant AR AD*^23^, with a helix-breaking mutation (L26P) in motif ^23^FQNLF^27^. Since, as shown above (Extended Data Fig. 3c,d), the helical character of regions or motifs rich in aromatic residues increases their propensity to engage in interactions we reasoned that this mutation would decrease the interaction of motif ^23^FQNLF^27^ with Tau-5*, thus allowing EPI-001 to more effectively target this sub-domain. In agreement with this hypothesis, EPI-001 induced higher CSPs in h2 and h3 of AR AD* than of AR AD (Extended Data Fig. 3e,f). Finally, we measured intermolecular PREs in construct allTau-5 to validate that high values of Δ*σ* are due to direct interactions induced by EPI-001: we produced constructs in which two of the three native Cys residues (C404, C448 and C518) were substituted by Ser, labelled the remaining Cys residue with a paramagnetic probe and measured its effect on the resonances of the allTau-5 enriched in ^15^N (Fig. 3c). In agreement with the CSP matrix, the regions of sequence of allTau-5 with aromatic and partially helical character interacted with one another, and addition of EPI-001 strengthened these interactions (Fig. 3c and Extended Data Fig. 3g,h).

Next we focused our attention on the sequence determinants of the selective interaction between EPI-001 and oligomeric AR AD. Since the interaction site of EPI-001 has a high helical propensity and a high density of aromatic residues (Extended Data Fig. 2e,f) we introduced amino acid substitutions in Tau-5* designed to investigate, independently, how these sequence features influence its interaction with EPI-001 (Fig. 4a). To alter helical propensity we introduced helix-breaking substitutions in helices h2 and h3 (A398P and L436P) and helix-stabilizing substitutions in helix h2 (a single substitution, G407A, and a triple substitution, termed AAA, including substitutions G394A, S395A and G407A) (Fig. 4a and Extended Data Fig. 4a). To alter aromatic character we replaced all aromatic residues with Ala (noAro) or, to achieve the same effect locally, substituted by Ala or Ser those in helix h2 (h2SA variant: Y393S, W397A, Y406S and H413A) or helix h3 (h3A variant: W433A, H434A, and F437A); we also replaced Cys 404 with Tyr (Fig. 4a), thus increasing aromatic character. We found that noAro has a higher helical propensity than WT, as expected given the high helical propensity of Ala residues, and that substitution C404Y had a negligible effect on the secondary structure of this construct (Extended Data Fig. 4a).

**Fig. 4.**
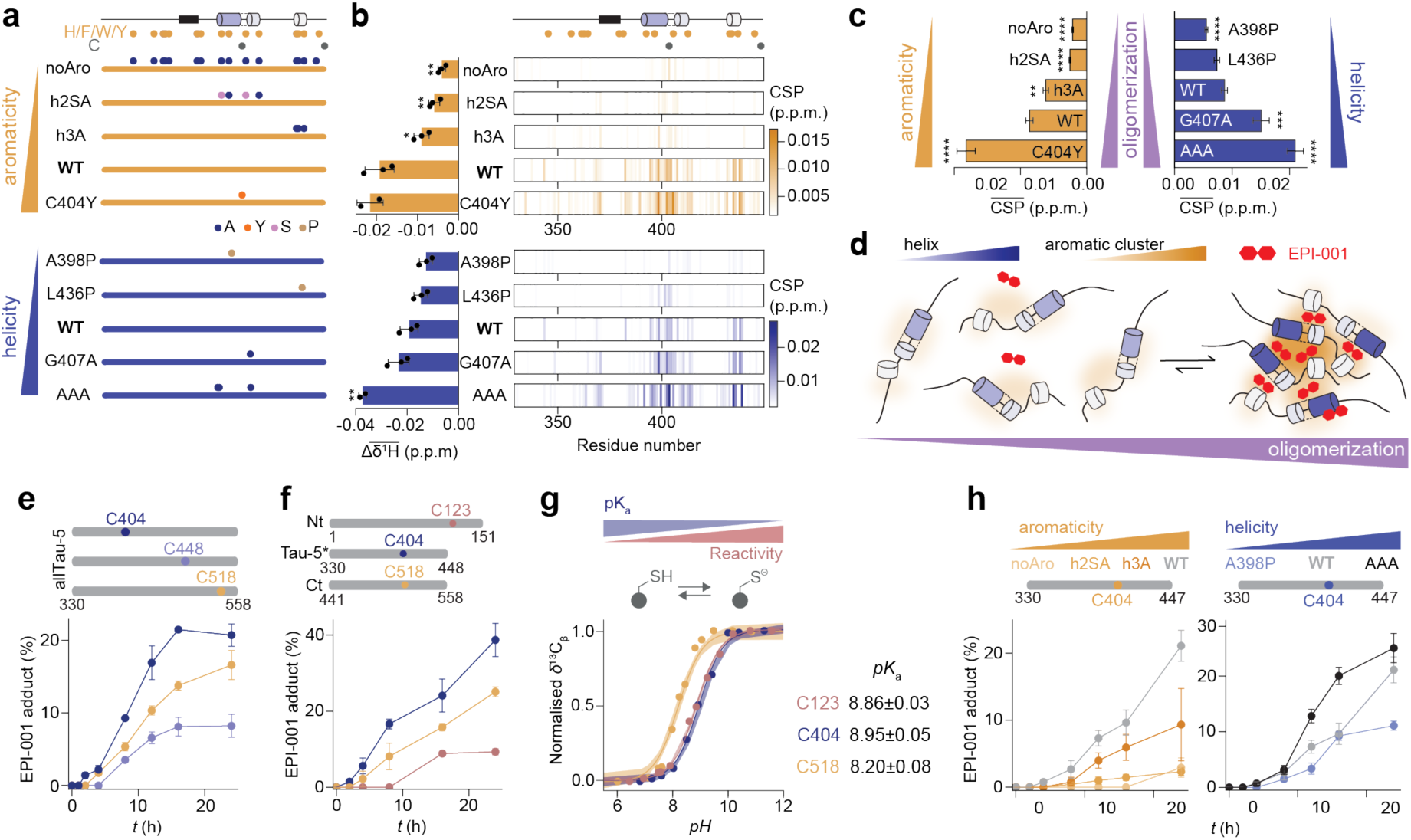
Tau-5* forms a transient EPI-001 interaction site upon oligomerization, induced by helical regions rich in aromatic residues. (**a**) Schematic of Tau-5* mutants. Orange and grey dots indicate the position of aromatic or cysteine residues, respectively. (**b**) *(Left)* Average shifts of the EPI-001 ^1^H signals induced by 1 molar equivalent of the corresponding Tau-5* mutant. *(Right)* Per-residue CSPs in the 2D ^1^H-^15^N NMR correlation spectra of Tau-5* upon addition of 1 molar equivalent of EPI-001. (**c**) Average protein CSPs induced by increasing the protein concentration from 25 to 400 µM in 2D ^1^H-^15^N NMR correlation spectra. (**d**) Scheme illustrating the preferential binding of EPI-001 to the oligomeric species that contain a higher density of aromatic patches and helical elements. (**e**) Percentage of covalent adduct as a function of time using intact mass spectrometry (MS, Extended Data Fig. 4g). Each allTau-5 mutant contains a single cysteine, while the remaining two were substituted to serines. (**f**) The percentage of covalent adduct formed in samples of AR AD fragments as a function of time using intact MS. Each AR AD fragment contains a single cysteine. (**g**) Cysteine *pK*_a_ values of AR AD fragments quantified by monitoring the *pH*-induced shifts in the cysteine ^13^C_β_ NMR signals. Errors indicate the standard error of *pK*_a_ estimation. (**h**) Percentage of covalent adduct, in sample od the Tau-5* mutants, as a function of time using intact MS. (**b**, **c**, **e, f, h**) Error bars denote standard deviations (*n* = 3).

We then investigated how these substitutions influence the interaction of Tau-5* with EPI-001, observing a clear correlation between the aromatic and helical characters of the variants and the size of the CSPs caused by their interaction with EPI-001 (Fig. 4b and Extended Data Fig. 4b). To investigate whether aromatic and helical character determine the ability of EPI-001 to interact with the AR AD because they are determinants of oligomerization we next measured the CSPs induced by increases in protein concentration (Extended Data Fig. 4c,d) in the absence of EPI-001. We obtained that Tau-5* oligomerization increases monotonically with aromatic character and helical propensity (Fig. 4c and Extended Data Fig. 4e), indicating that oligomeric Tau-5* is stabilized by interactions between aromatic side chains in regions of sequence rich in secondary structure and that the selective interaction between EPI-001 and Tau-5* relies on the oligomerization of its target (Fig. 4d).

It has been proposed that EPI-001 is a covalent AR inhibitor that reacts with nucleophilic side chains of Cys residues in the AR AD (Extended Data Fig. 4f)^11^. To determine whether its interaction with the target influences the rate of covalent attachment to Cys residues, we measured the reaction rates using three variants of the allTau-5 construct, which contains three such residues (C404, C448, and C518) by using mass spectrometry (MS, Extended Data Fig. 4g). In each variant, two Cys residues were mutated to Ser, leaving a single Cys available for reaction. We found that the reaction was markedly faster with Cys 404 (Fig. 4e), located in Tau-5* region which harbors the site of interaction of EPI-001. To further confirm that Cys 404 exhibits the highest reactivity, we examined the covalent modification rate of AR AD fragments—Nt, Tau-5*, and Ct—each containing only one Cys residue (C123, C404, or C518, respectively). Additional Cys residues were either mutated to Ser (C129S in Nt) or deleted (C448 in Tau-5* and Ct). Consistent with the allTau-5 variant analysis, Cys 404 reacts faster (Fig. 4f). This difference in reactivity is not due to differences of nucleophilic character as determined indirectly by measuring the *pK*_a_ of the thiol group of the difference Cys residues ^29–31^ (Fig. 4g). Finally, we investigated how the reactivity of Cys 404 is influenced by the strength of the reversible interaction between EPI-001 and Tau-5*. We found that reactivity correlates directly with the helical and aromatic character of the construct (Fig. 4h), suggesting that the enhanced reactivity of Cys 404 results from the prolonged residence time of EPI-001 at its interaction site within Tau-5*. In addition, since the detection by MS of the adduct provides direct evidence for the reaction between Tau-5* and EPI-001 this results confirms that CSPs reliably report on the strength of the reversible interaction between these EPI-001 and its target.

### Effects of EPI-001 partitioning in NR AD condensates

Several independent studies have established that the activity of AR as a transcription factor relies on its capacity to condense upon DNA binding^23,32–34^ and thus phenotypic screens based on the high-throughput analysis of AR condensation dynamics, rather than on the measurement of transcriptional activity, can be used to identify new AR inhibitors^32^. More generally, since biomolecular condensates represent liquid phases distinct from the solutions surrounding them it has been suggested that drug-like small molecules may selectively partition in them without necessarily engaging in stereospecific interactions, a process that could facilitate target engagement and therefore be exploited for drug discovery and development^35,36^. This concept has recently been tested both *in vitro* and in cells, revealing that the physico chemical properties of condensates are indeed related to those of the drug-like small molecules that partition in them^37,38^. As we previously showed^23^, the aromatic character of the AR AD is key for condensation likely due to the ability of aromatic residues to engage in π-π interactions. These interactions can be homotypic, with the ADs of other AR molecules, or heterotypic, with other components of transcriptional condensates such as RNAPII which, like the AR AD, is enriched in Tyr residues in its C-terminal domain, that is intrinsically disordered^23,32–34^. We also showed that EPI-001 partitions in the condensates formed by the AR AD and that modifying the structure of this drug-like small molecule to increase its affinity for the AR AD as well as its partitioning in AR AD condensates increase its potency as AR inhibitor^23^.

To determine whether selective partitioning contributes to selectivity, we measured the partition coefficient of EPI-001 in the condensates formed by the selectivity panel *in vitro*. We found that the GR AD does not form droplets *in vitro* (Extended Data Fig. 5a) and that EPI-001, bearing two aromatic rings, hardly partitions in PR and MR ADs droplets (Fig.5a). By contrast, EPI-001 partitions similarly in the condensates formed by the ERα AD, which is the AD most enriched in aromatic residues, as well as those formed by the AR AD that has a lower aromatic character (Fig. 1c) but harbors the interaction site of EPI-001 (Fig.5a). These results suggest that small molecules bearing aromatic rings partition in condensates formed by sequences rich in aromatic residues, in line with recent findings^23,37,38^, but that stereospecific interactions can also contribute to partitioning. We next studied whether EPI-001 partitioning into NR AD droplets *in vitro* correlates with the behavior in cells. For this we used a cell-based condensate tethering system, in which the NR ADs are expressed as a fusion protein with the DNA-binding domain of the Lac repressor (LacI) and Cyan Fluorescence Protein (CFP) in cells containing an integrated array of LacO binding sites (CFP-LacI-IDP)^39^. In this experimental set-up the enrichment of a second, fluorescently tagged protein can be visualised and quantified in the tethered condensate^34,40–42^ (Fig. 5b).

**Fig. 5.**
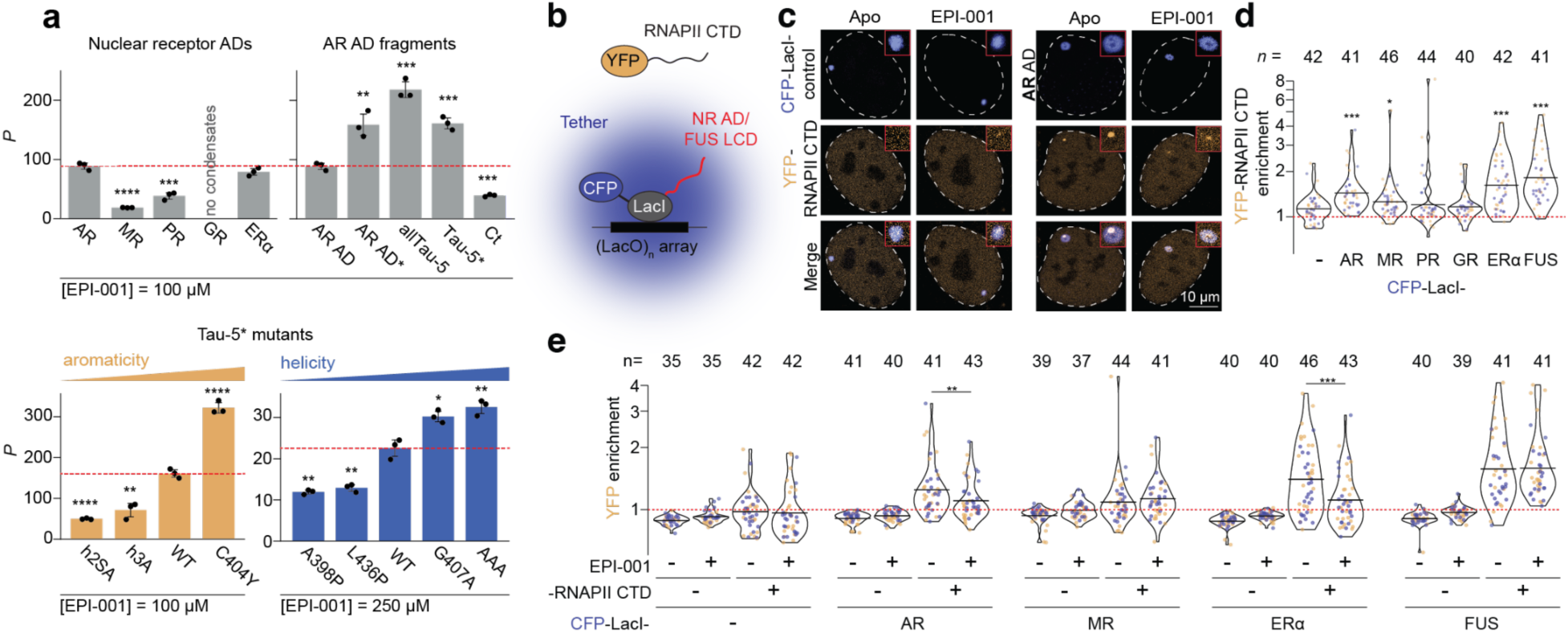
EPI-001 partitioning reduces the ability of the NR condensates to recruit RNAP II CTD in live cells. (**a**) EPI-001 partition coefficients into condensates of NR ADs, AR AD fragments and Tau-5* mutants. (**b**) Illustration of the condensate tethering experiment. (**c**) Exemplary images of cells co-transfected with YFP-RNAPII CTD and CFP-LacI-AR AD expression vectors. Cells were treated either in the absence (Apo) or presence of 25 µM EPI-001. Each image depicts a single nucleus, delineated by a white dashed line. Within the red squares, a zoomed-in version of the CFP focus is displayed. Notably, the RNAPII CTD signal does not entirely overlap with the tether, consistent with known characteristics of RNAPII CTD recruitment in this assay^40,41^. (**d**) Quantitative analysis of YFP-RNAPII CTD enrichment in NR AD and FUS LCD tether foci. Each data point represents a single tether/cell, with the number of tethers acquired for each condition indicated above the plot. Data are compiled from two independent transfections (orange and blue). (**e**) Quantification of YFP(-) or YFP-RNAPII CTD(+) enrichment in various tethers. Enrichment levels of RNAPII CTD in DMSO(-) and EPI-001(+)-treated samples were normalised to RNAPII CTD co-transfected with empty LacI-CFP under corresponding conditions. Data originate from two independent transfections, distinguished by colour (orange and blue).

We measured the recruitment of RNAPII C-terminal domain (CTD), a key client in transcriptional condensates^43^, as a proxy of co-partitioning driven by aromatic residues^23,44,45^. RNAPII CTD was fused with Yellow Fluorescence Protein (YFP) into lacO-induced condensates formed by the NR ADs in the condensate tethering system (Fig. 5c and Extended Data Fig. 5b,c). The FUS Low-Complexity Domain (LCD) was included as a positive control, as RNAPII CTD is known to partition into FUS LCD condensates^46,47^. RNAPII CTD was significantly enriched in condensates formed by AR and ERα ADs and to a lesser extent in those formed by MR AD but showed no significant enrichment in GR and PR AD condensates (Fig. 5d and Extended Data Fig. 5d). Then, to test the effect of EPI-001 on the recruitment of RNAPII CTD into condensates, we measured RNAPII CTD levels in the presence of the compound for the condensates that were enriched with RNAP II CTD: AR AD, MR AD, ERɑ AD, and FUS LCD. In good agreement with *in vitro* results (Fig. 5a), we observed a significant reduction in the enrichment of RNAPII CTD in both AR AD and ERɑ AD condensates upon treatment with the small molecule, but not in MR AD and FUS LCD condensates (Fig. 5e and Extended Data Fig. 5b-d). These findings indicate that EPI-001 partitions into transcriptional condensates and that interactions between aromatic rings remain relevant in a cellular environment.

We next investigated the role of stereospecific interactions in partitioning. For this we measured the partition coefficient of EPI-001 in the droplets formed by AR AD fragments and variants that interact with this small molecule to different extents (Fig. 2d and Extended Data Fig. 2g). First, we used constructs allTau-5, Tau-5* and Ct (Fig. 5a and Extended Data Fig. 5a). EPI-001 partitioned better in those containing the interaction site of EPI-001, Tau-5* and allTau-5, than in Ct, despite it being rich in aromatic residues (Fig. 5a and Extended Data Fig. 2e). EPI-001 partitioned better in Tau-5* and allTau-5 than in full-length AR AD, that possesses the motif ^23^FQNLF^27^: since this motif competes with EPI-001 for Tau-5* (Extended Data Fig. 3e,f) the partition coefficient of this small molecule is higher in AR AD* droplets, where a helix-breaking substitution (L26P) weakens the interaction between ^23^FQNLF^27^ and Tau-5*. We also measured the partition coefficient of EPI-001 in Tau-5* variants (Fig. 5a and Extended Data Fig. 5a): we found that substitutions in Tau-5* that reduce the interaction between EPI-001 and Tau-5* and, therefore, the reactivity of Cys 404—by decreasing the aromatic character or helical propensity of the site of interaction—decreased EPI-001 partitioning (Fig. 5a). Collectively, our results indicate that the interaction between EPI-001 and the AR AD has a strong influence on its partitioning in the AR AD droplets.

Finally, to study how EPI-001 partitioning influences the properties of the AR AD droplets *in vitro*, we used three different constructs bearing the interaction site of this small molecule: AR AD, allTau-5 and Tau-5* (Fig. 2c). In all cases we found that EPI-001 increased the number of arrested fusion events (Fig. 6a,b and Extended Data Fig. 6a), indicating a reduced dynamic character of the droplets, which we confirmed by using fluorescence recovery after photobleaching (FRAP) experiments (Fig. 6c).

**Fig. 6.**
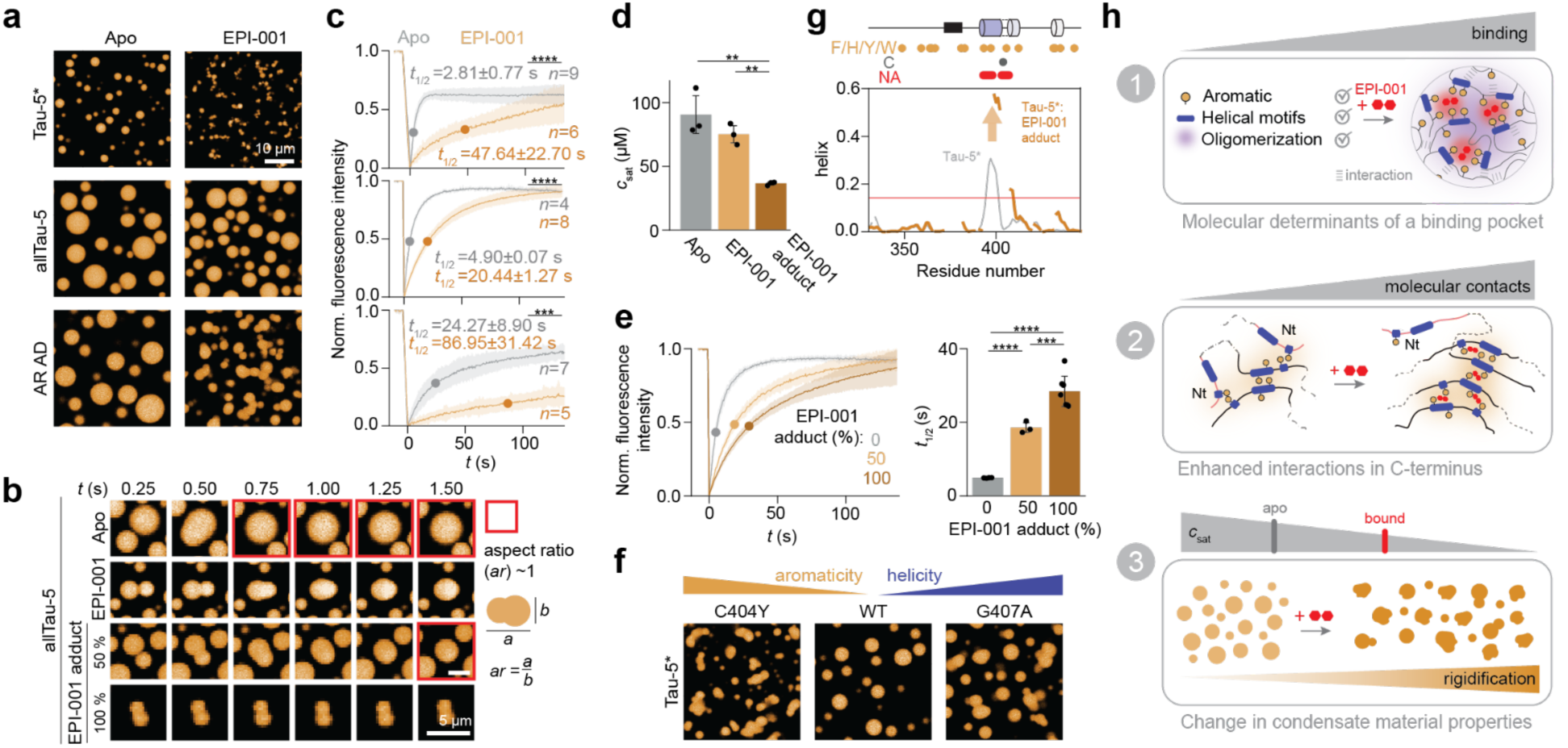
EPI-001 binding modulates AR AD condensates and induces their rigidification. (**a**) Fluorescence microscopy images showing *in vitro* reconstituted condensates formed by the AR AD constructs that contain the binding site (Tau-5*, allTau-5 and AR AD) both in the absence and presence of EPI-001. (**b**) Fusion events monitored at different time points of allTau-5 droplets. Both the reversible interaction and covalent adduct forms of EPI-001 were measured. Spherical fused droplets are highlighted in red. (**c**) FRAP experiment of the AR AD fragment droplets. (**d**) Saturation concentration (*c*_sat_) measurements of allTau-5 in the presence and absence of EPI-001 (ratio 1:1), and allTau-5*:EPI-001 adduct (100 %). Reduced *c*_sat_ indicates an enhancement of the condensation by the ligand. (**e**) FRAP experiments of allTau-5 condensates at different percentages of covalent adduct. (**f**) NMR-derived helical content of Tau-5* and Tau-5*:EPI-001 adduct. Orange and grey circles indicate the positions of aromatic and cysteine residues, respectively. Red circles indicate not assignable residues due to line broadening. The red line indicates the uncertainty of the *δ*2D algorithm (5%)^51^. (**g**) Fluorescence microscopy images of Tau-5* mutants. (**h**) (*1*) Schematic illustration depicting the molecular basis of the transient binding site formed by AR AD molecules. (*2*) Diagram showing how EPI-001 enhances interactions between AR AD molecules, rewiring the network of interactions and leading to increased oligomerization. (*3*) The drug effect on AR AD condensates: enhanced phase separation and rigidified condensates. (**d, e**) Error bars indicate standard deviations (*n* => 3).

Since EPI-001 may inhibit the AR in part by reacting with Cys residues in the AR AD, we analysed how the covalent attachment of EPI-001 influences the properties of the condensates (Fig. 6b). We found that the saturation concentration (*c*_sat_) and cloud temperature (*T*_c_) decrease upon covalent attachment, thus enhancing condensation (Fig. 6d and Extended Data Fig. 6b), as well as decrease the dynamic character of the droplets as monitored by FRAP experiments (Fig. 6e). Previous studies have shown that the clustering of aromatic residues in IDRs that undergo phase transitions can lead to aggregation^48^. In agreement with this notion we found that the introduction of an additional aromatic residue by substitution of Cys 404 (C404Y) promotes oligomerization (Fig. 3f) and reduces the dynamic character of droplets (Fig. 6f and Extended Data Fig. 6c), as does a variant that increases the helical propensity of h2 (G407A, Fig. 6f and Extended Data Fig. 6c). To determine whether this also occurs upon covalent attachment of EPI-001 to the AR AD, a reaction that increases the aromatic character of the sequence by introducing two aromatic rings, we used solution NMR. We found that the covalent attachment of EPI-001 increases the helical propensity of helix h2 two-fold (Fig. 6g and Extended Data Fig. 6d). Together, these findings suggest that increased aromaticity and helicity in h2, an aromatic-rich region, contribute to condensate hardening, likely via enhanced aromatic clustering mediated by this secondary structure element.

## Discussion

Targeting IDRs of proteins remains a major challenge in drug discovery, primarily due to their lack of well-defined binding sites required for small-molecule interactions. Unlike globular proteins, which present pockets for ligand binding, IDRs exhibit conformational flexibility and dynamic interactions, making traditional drug-targeting strategies less effective^2^. Despite this difficulty, our study shows that drug-like small molecules can selectively interact with IDRs by exploiting transiently structured conformations.

We investigated the interaction between EPI-001 and the activation domain of the androgen receptor. EPI-001 is particularly relevant as it represents one of the first small molecules targeting an IDR to reach clinical trials, having shown efficacy in preclinical models^10,11^. Our motivation for this study stemmed from the need to understand the molecular basis of its selectivity and to explore generalizable principles for targeting IDRs with small molecules.

Our findings reveal that EPI-001 selectively binds an oligomeric form of the AR AD. This oligomeric state is more structured than the monomeric form and is stabilized by interactions involving aromatic residues in regions with helical propensity (Fig. 6h). Unlike other NR ADs, AR AD contains helical elements and aromatic residues that drive its oligomerization, likely explaining its selective binding to EPI-001 (Extended Data Fig. 1i). This suggests that IDR oligomerization can lead to the transient formation of structured conformations that provide opportunities for small-molecule targeting of this class of proteins. The selective stabilization of these transiently structured states represents thus a novel avenue for drug discovery against IDRs, conceptually related to the exploitation of cryptic binding pockets for drug discovery against globular targets^49^.

This concept is particularly relevant for transcription factors, as increasing evidence suggests that the propensity of their IDRs to self-assemble plays a key role in their biological activity^50^. Our results indicate that drug-like small molecules can exploit this to achieve selective inhibition. In the case of EPI-001, its interaction with the AR AD not only stabilizes a specific oligomeric conformation but also rewires the conformational ensemble of the target, thereby altering the physical properties of its phase-separated (Fig. 6h) state and ultimately reducing its transcriptional activity.

Our study also highlights the importance of both physicochemical properties^37,38^ and stereospecific interactions in rationalizing the partitioning of drug-like small molecules in biomolecular condensates: EPI-001 selectively partitions into condensates formed by the AR AD due to its aromatic character as well as to specific molecular interactions with the target. These results indicate that optimizing both the physicochemical properties of small molecules and their interactions with their targets will be important in the discovery of drugs targeting proteins forming biomolecular condensates, as we have recently shown for the AR AD^23^.

In summary, our study shows that IDR oligomerization can provide opportunities for selectively targeting this class of proteins with drug-like small molecules. The insights gained here pave the way for rational design strategies aimed at developing therapeutics against transcription factors and other IDR-containing proteins. In addition, they emphasize the importance of considering condensation and oligomerization as key factors in the development of inhibitors for intrinsically disordered targets.

## Materials and Methods

### Sequence analysis

NR protein sequences (UNIPROT ID: P10275 (AR), P08235 (MR), P06401 (PR), P04150 (GR) and P03372 (ERɑ)) where used to calculate the sequence identity for the ADs and the folded domains, the conserved N-terminal cysteine of the DBD (C-V/A-I/V-C motif) was arbitrarily set to classify between the AD and the DBD-LBD. In this study, ERɑ AD was taken as prototypical of the ER nature. As control, the sequence properties were also calculated for the IDR regions of TDP-43 (residues 262-414. UniProt ID: Q13148), FUS (residues 1-270. UniProt ID: P35637), EWS low-complexity domain (residues 1-264. UniProt ID: B1PRL2), Nup98 (residues 1-741. UniProt ID: P52948), hnRNAP1 (residues 186-372, UniProt ID: P09651), and CPEB4 (residues 1-448, UniProt ID:Q17RY0).

The sequence properties of the intrinsically disordered NR ADs were calculated using the CIDER package^18^. The calculated parameters were the mean hydropathy using the Kyte-Doolittle scale^57^; Fraction of disorder promoting residues in the sequence; *κ* and *Ω* values describe the segregation of charged and proline residues as previously described^48,54^; FCR, fraction of charged residue; and the NCPR, net charge per residues. The aromatic clustering (*Ω*_aro_) was calculated by making a sequence length correction of a previously proposed parameter^56^. This metric computes the average inverse distance among aromatic residues and normalizes it by the sequence length by calculating the *Ω*_aro_ value for a maximum clustered sequence (*Ω*_aro,clust._) and an ideally spaced aromatic sequence (*Ω*_aro, ideal._):

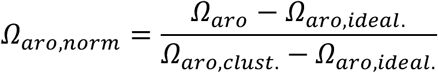

The sequence patterning plots shown in Extended Data Fig. 1b, were calculated by measuring a rolling mean average of 30 residue windows. The hydropathy plot took into account the Kyte-Doolittle scale^57^. For the charge plot, arginine and lysine residues had associated a +1 value, aspartate and glutamate residues a -1 value at *pH* 7.4.

To quantify the sequence patterning we calculated *z*-scores using the NARDINI program^16,53^. The *z*-score reflects on the degree of blockiness of groups of residues compared to 10^6^ randomly generated sequences with the same composition. Residues are grouped into the following eight types: polar≡(Q,S,H,T,C,N), hydrophobic≡(I,L,M,V), positive≡(K,R), negative≡(D,E), aromatic≡(F,Y,W), alanine≡A, proline≡P, and glycine≡G. Considering all pairs of residue types leads to 36 patterning features, where 6 correspond to *Ω* parameters – same type segregation – whereas the other 30 are the patterning of different class comparisons or *δ*-parameter. The positive *z*-scores imply the patterning of the two residue types is more blocky than random, whereas negative *z*-scores imply the patterning is more well-mixed.

To visualise the disordered region of the activation domain of AR (Fig. 1a), the DODO package was used (https://github.com/ryanemenecker/dodo).

### Protein sequences

The sequences of proteins used for the *in vitro* experiments for the Article are provided in Table S1 (Supplementary Information).

### Protein expression

Recombinant expression of non-isotopically labelled proteins were produced in Lysogeny Broth (LB), whereas isotopically labelled (^15^N or double-labelled ^15^N and ^13^C) proteins were expressed in minimal M9 media supplemented with ^15^NH_4_Cl and, if needed, ^13^C-glucose. Cell cultures at *OD*_600_ = 0.6 were induced with 1 mM isopropyl 𝛽-D-1 thiogalactopyranoside (IPTG) and incubated under specific conditions for expression.

Expression trials were performed for most of the protein constructs used in this study. Depending on the protein, different expression vectors, bacterial cell strains or expression conditions were chosen. The different AD of the NR family: AR (1-558 aa, with 21Q and 24G polyQ and polyG tracts at positions 58-78 and 449-472, respectively), AR* (L26P)^23^, AR C404 (all cysteines deleted, except C404), MR (1-602 aa), PR (1-566 aa), GR (1-420 aa), ERɑ (1-184 aa); Ct constructs (441-558 aa): Ct, Ct 4G (containing 4G in polyG tract), Ct C518 (containing 4G in polyG tract and the C448 is deleted); and Tau-5* constructs (330-448 aa): Tau-5* and it is mutants

- A398P+L436P, C404Y, A398P, L436P, G407A, C404S and C404 (lacking the last residue C448)
- were recombinantly produced in *E. coli* Rosetta (DE3) cells. Upon induction with IPTG, the expression was carried out in LB or M9 media at 25°C overnight under agitation. noAro (where all the aromatic residues were substituted to alanines), AAA (G394A+S395A+G407A), A398P+T435P and T435P mutants of Tau-5* were produced in *E. coli* BL21 (DE3) cells (New England Biolabs #C2527) at 25°C overnight in both LB and M9 media. While h2SA (Y393S+W397A+Y406S+H413A) and h3A (W433A+H434A+F437A) mutants of Tau-5* were produced in *E. coli* BL21 (DE3) LysY (New England Biolabs #C3010I) cells at 25 °C overnight for unlabelled LB media and 37°C for 3 hours for ^15^N/^13^C M9 media. In all cases, the protein constructs were cloned in pDEST™17 vector (ThermoFischer Scientific).

The allTau-5 construct (330-558 aa, polyG with 4G, to prevent protein aggregation^58^) and allTau-5 24G (polyG with 24G), allTau-5 mutants CtoS (C404S+C448S+C518S), CtoS PP (C404S+C448S+C518S+A398P+L436P), and mutants containing only one cysteine C404, C448, and C518 (with the other two cysteines mutated to serines), were produced in *E. coli* BL21 (DE3) LysY cells (New England Biolabs #C3010I) transformed with pET-30a(+) plasmids (Novagen) at 25°C overnight in LB and M9 media.

For non-isotopically labelled Nt constructs (1-151 aa): Nt, Nt L26P, and Nt C123 (with C129S substitution), *E. coli* BL21 (DE3) LysY cells (New England Biolabs #C3010I) were transformed with a pET-50b(+)plasmid (Novagen) with the solubility tag of NusA preceding the protein of interest cloned. Protein was expressed overnight at 37 °C upon induction with IPTG. Isotopically labelled Nt C123 was produced under the same conditions but expressed at 25 °C. Isotopically labelled Nt and Nt L26P were produced in *E. coli* Rosetta (DE3) and BL21 (DE3) cells, respectively, transformed with pDEST™17 vector (ThermoFischer Scientific) containing maltose binding protein (MBP) tag and expressed in M9 media overnight at 25 °C. The use of solubility tags such as NusA and MBP prevented Nt aggregation^25^. Notably, the pDEST™17 vector with the MBP tag displayed higher yields in protein production in M9 media compared to pET-50b(+) with NusA.

Every plasmid was engineered to contain a His-tag following the solubility tag (for Nt constructs) and either a His-3C or a TEV (Ct and Nt variants) cleavage site upstream of the encoded protein sequences.

### Protein purification

The purification was performed as previously reported^23–25^. Briefly, for all protein constructs except Nt, cells were harvested by centrifugation and resuspended in phosphate-buffered saline (PBS). Then the cells underwent two rounds of sonication for 7 minutes each with a 5-second on-and-off pulse. Following centrifugation, the supernatants were discarded, and the pellets were washed twice using a Wash buffer (PBS, 1% Triton, 500 mM NaCl, 1 mM dithiothreitol (DTT), *pH* 7.8). 5 mM MgSO_4_ and 130 µM CaCl_2_ were added in the first wash. Insoluble inclusion bodies were collected via centrifugation and subsequently solubilized overnight at room temperature in a binding buffer (20 mM Tris, 500 mM NaCl, 5 mM imidazole, 8 M Urea, 0.05% w/v NaN_3_, 1 mM DTT, *pH* 7.8). The solubilized inclusion bodies underwent centrifugation, filtration of the supernatant, and were applied to a HisTrap HP column (Cytiva) at room temperature and eluted with a gradient of 500 mM imidazole in the binding buffer. Eluted fractions underwent two rounds of dialysis, of 3 hours and 16 hours, using a buffer (50 mM Tris, 1 mM DTT, *pH* 8.0). His-tag cleavage was performed using either TEV or 3C proteases during the second dialysis step. The addition of 0.5 mM EDTA in the dialysis buffer for TEV protease was needed. After dialysis, urea was added to the sample to reach a final concentration of 8M a and a second HisTrap HP column (Cytiva) was performed at room temperature to collect the cleaved protein in the flow-through. Concentration was achieved using 3 kDa or 10 kDa Amicon® concentrators (Merck) depending on the molecular weight of the respective protein. The samples were aliquoted and stored at -80°C. Notably, the allTau-5 and Ct constructs were better purified by adjusting the urea concentration to 6 M and adding 2 M GuHCl in the binding buffer.

For the Nt constructs, cells were harvested via centrifugation and resuspended in a Core buffer (20 mM NaH_2_PO_4_, 500 mM NaCl, 5% v/v Glycerol, 0.05% w/v NaN_3_, 1 mM DTT, *pH* 8). Lysozyme powder was added to facilitate cell lysis at 1.5 mg/mL concentration, followed by incubation for 1 hour at 4 °C on a spinning wheel. The lysed cells were then subjected to sonication for 20 minutes with a pulse of 5 seconds on and 10 seconds off. Upon centrifugation, the supernatant was filtered and loaded onto a HisTrap HP (Cytiva) column at 4 °C. Fractions containing the protein of interest were eluted from the column using a gradient of 500 mM imidazole in the Core buffer. The eluted fractions were concentrated using 10 kDa Amicon® concentrators (Merck) and injected on a size exclusion chromatography (SEC) using an HiLoad superdex 200 pg 26/600 column (Cytiva). The purest fractions were collected and mixed with either TEV or 3C protease. Subsequently, dialysis was conducted in a Core dialysis buffer (50 mM NaH_2_PO_4_, 100 mM NaCl, 1 mM DTT, *pH* 8, with the addition of 0.5 mM EDTA for TEV) at 4 °C overnight. After dialysis, urea was added to the sample to reach a final concentration of 8M and a second HisTrap HP column (Cytiva) was performed at room temperature to isolate the cleaved protein which is present in the flow-through. This protein was concentrated using 3 kDa concentrators, aliquoted, and stored at -80 °C.

### Small molecule preparation

EPI-001 was obtained commercially from Sigma-Aldrich (SML1844) and used in the experiments.

### Protein sample preparation

Stored protein aliquots at -80 °C were thawed and subjected to SEC using HiLoad superdex 75 or 200 pg columns (Cytiva) in a final buffer (20 mM sodium phosphate, 1 mM tris(2- carboxyethyl)phosphine (TCEP), 0.05% w/v NaN_3_ at pH 7.4), maintained at 4 °C unless specified otherwise. After the SEC, the fractions containing the pure protein were concentrated using either a 3 kDa or 10 kDa Amicon® concentrator (Merck) depending on the protein construct size. Subsequently, samples underwent centrifugation to remove aggregate traces, and their concentrations were determined using a microvolume spectrophotometer (NanoDrop, Thermo Fisher Scientific) microvolume spectrophotometer at 280 nm, except for noAro samples, where determination via absorbance was not feasible due to the absence of aromatic residues, thus requiring the use of an High performance liquid chromatography (HPLC) system. The absolute concentration of an noAro sample was initially determined using amino acid analysis. Subsequently, various dilutions were prepared to build a calibration curve using an HPLC system. This calibration curve was constructed by measuring the protein peak integration, detected at approximately 49% of ACN:H_2_O, across different concentrations. This provided a direct relationship between the protein concentration and the corresponding peak integral. The final samples were prepared on ice.

The samples involving the small molecule EPI-001, including their Apo versions, contained 0.5% DMSO unless otherwise specified. EPI-001 was added from 50 or 100 mM DMSO or DMSO-d_6_ (for NMR experiments) stocks.

### Spin-labelling for PREs measurement

Purified allTau-5 Cys-mutants —C404, C448, and C518— and AR C404 were reduced by incubating with 5 mM DTT, after which the solution was buffer-exchanged into a TCEP-free final buffer (see Sample preparation) by passing the protein solution through a size-exclusion column HiLoad superdex 75 or 200 pg 13/300 (Cytiva). Proteins were then incubated with 5 molar equivalents of S-(1-oxyl-2,2,5,5-tetramethyl-2,5-dihydro-1H-pyrrol-3-yl)methyl methanesulfonothioate (MTSL) (Toronto Research Chemicals), which was added from aDMSO stock, and allowed to react overnight with agitation at 37 °C and pH 8. Unreacted MTSL and disulfide-bridged protein dimers were removed by SEC using a HiLoad superdex 75 or 200 pg 16/600 column (Cytiva). Nearly 100% cysteine modification was confirmed using intact MS.

### Cysteine thiol group blocking by iodoacetamide

Iodoacetamide (Merck) DMSO stocks, freshly prepared at 1M, were used to block cysteines of Tau-5* C404S (to create 1Tau-5*) and AR AD C404. First, proteins were reduced by incubating with 5 mM DTT, after which the solution was buffer-exchanged into a TCEP-free final buffer (see Sample preparation) by passing the protein solution through a size-exclusion column HiLoad Superdex 75 or 200 pg 13/300 (Cytiva). Then, protein samples were incubated with 10 molar equivalent of iodoacetamide, at pH 8 and 37 °C for 1 hour. Unreacted iodoacetamide and disulfide-bridged protein dimers were removed by SEC using a HiLoad Superdex 75 or 200 pg 16/600 column (Cytiva) with the TCEP-free final buffer.

### Generation of 2Tau-5*

The Tau-5* C404S sample was buffer-exchanged into a TCEP-free final buffer at pH 8 (see Sample Preparation) by passing the protein solution through a size-exclusion column, HiLoad Superdex 75 13/300 (Cytiva). Some dimer formation via disulfide bridge between C448 (2Tau- 5*) occurred on the column. Monomeric peak of Tau-5* C404S was collected and incubated at pH 8 and 37 °C for 48 hours. Monomeric Tau-5* C404S and 2Tau-5* were then separated using a size-exclusion column, HiLoad Superdex 75 16/600 (Cytiva), with the TCEP-free final buffer.

### Generation of the EPI-001 protein adduct

The allTau-5:EPI-001 adduct was prepared by incubating allTau-5 with EPI-001 at a 1:5 molar ratio in the final buffer at *pH* 8 and 37 °C for 72 hours. Afterwards, a SEC using an HiLoad superdex 75 pg 26/600 column (Cytiva) was conducted to eliminate formed multimers and free EPI-001. The efficacy of the reaction was evaluated through intact MS (Supplementary Information: Fig. S2). For ^15^N, ^13^C-double labelled Tau-5*:EPI-001 adduct, Tau-5* C404, which contains one cysteine per molecule (i.e. C448 was deleted), was incubated with EPI-001 at a 1:10 molar ratio in the final buffer at *pH* 8 and 37 °C for 96 hours. Separation of the adduct from the protein and free compound was achieved using a Jupiter C4 semiprep column (Phenomenex) connected to an Agilent Technologies 1200 HPLC instrument. Mobile phases consisting of H_2_O and ACN:H_2_O (9:1) with 0.1% v/v trifluoroacetic acid (TFA) were utilised. The elution of Tau-5*:EPI-001 occurred with 45% v/v ACN:H_2_O. The fraction containing the adduct was collected and lyophilized, followed by redissolution in the final buffer with the *pH* readjusted to 7.4.

The concentrations of both adducts were ascertained by absorbance at 280 nm using NanoDrop (Thermo Fisher Scientific) from extrapolation of a calibration curve. A known concentration sample underwent amino acid analysis to derive its exact concentration, and a dilution series of the unknown sample concentration amount was subsequently measured with the NanoDrop to establish the absorbance dependence on concentration.

### Amino acid analysis for protein quantification

The protein samples were hydrolyzed in glass tubes with HCl 6 M (0.1-1% v/v phenol) for 24 h at 110 °C. The solvent was evaporated and the remaining powder redissolved in water and filtered. An aliquot of the filtered sample containing the amino acids (AA) was derivatized with 6- Aminoquinolyl-N-hydroxysuccinimidyl carbamate (AQC) following the indication of the AccQ-Tag method (Waters) to obtain the corresponding AQC analogs (AQC-AA). The derivatized AQC- AA were injected in a Nova-Pak C18 (4 µm) HPLC column (Waters) connected to a WATERS 600 HPLC system with a detector WATERS 2487. The amount of AQC-AA was followed by measuring the absorbance at 254 nm. Concentration of amino acids was calculated by an internal standard method. A known amount of aminobutyric acid (AABA), as internal standard, was added to the sample and analyte amount is calculated using area responses of analytes and internal standard. Working amino acid standard solutions were prepared by dilution of commercial 2.5 mM stock (Amino Acids Mix Solution 79248 Sigma-Aldrich). Internal standard solutions (2.5 mM) are prepared using Norleucine and AABA (Sigma-Aldrich).

### NMR experiments

All NMR experiments were recorded at 5 °C using Bruker Ascend Evo 1 GHz, Bruker Avance Neo 800 MHz and Bruker Avance III 600 MHz spectrometers equipped with TCI cryoprobes. Samples, unless otherwise specified, were measured in 3 mm NMR tubes containing the final buffer (20 mM sodium phosphate buffer, *pH* 7.4, 1 mM TCEP, 0.05% w/v NaN_3_) 10 µM sodium 3-(trimetylsilyl)propane-1-sulfonate (DSS) for chemical shift referencing, and 10% v/v D_2_O. All 3D triple-resonance experiments were acquired with non-uniform sampling applied to both indirect dimensions.

Data processing encompassed qMDD^59^ for non-uniformly sampled data reconstruction and NMRPipe^60^. Subsequent analysis was carried out using CcpNMR Analysis^61^.

### Assignments

Backbone assignment of ^15^N, ^13^C-double labelled ERɑ AD (150 µM) and GR AD (280 µM) was accomplished using the BEST-TROSY versions^62^ of 3D HNCO, HNCA, HNCACB, and HNcoCACB experiments in combination with 3D HNcaCO and hNcaNNH experiments^63^.

For backbone assignment of PR AD, a 200 µM ^15^N,^13^C-double labelled sample prepared in the final buffer *pH* 5.8 and 8% v/v D_2_O was used. BEST-TROSY versions of 3D HNCO, HNcaCO, HNCA, HNcoCA, HNCACB, and HNcoCACB, in combination with 3D HNcaCO and hNcocaNNH and 4D HNCACO were recorded.

For MR AD, two 200 µM samples were used, one in the absence and the other in the presence of 2 M urea, containing 2% v/v D_2_O. For each sample, BEST-TROSY versions of 3D HNCO, HNCA, HNCACB, and HNcoCACB experiments in combination with 3D HNcaCO and hNcaNNH experiments were acquired.

Backbone assignments for Nt and Ct were previously reported (BMRB ID: 25607 and 51476, respectively)^23,25^. For the ^15^N,^13^C-double labelled allTau-5, allTau-5 CtoS, and allTau-5 CtoS PP at 150 µM, assignments were accomplished through a series of BEST-TROSY versions of 3D experiments, including HNCO, HNCA, HNCACB, HNcoCACB, and HNcaCO. In addition, the 3D hNcaNNH experiment was measured for allTau-5. AllTau-5 24G assignments were derived from allTau-5 and confirmed using HNCA and HNCO experiments.

Backbone assignment of ^15^N,^13^C-double labelled Nt L26P at 150 µM was achieved using 3D HNCA, HNCO, HNcaCO, and HNCACB experiments. Assignments for 150 µM Tau-5* mutants (noAro, C404Y, A398P, G407A, AAA), 120 µM Tau-5* C404, 12 µM Tau-5*:EPI-001 adducts, and 0.31 mg/mL 1Tau-5* and 2Tau-5* were obtained through BEST-TROSY versions of 3D HNCO and HNCA experiments. Additional spectra recorded were HNcaCO for noAro, A398P, and AAA; HNCACB for noAro and AAA; and HNcoCACB for noAro, G407A, A398P, and Tau- 5*:EPI-001 adduct.

Assignments for h2SA and h3A were derived from Tau-5* and noAro assignments, while those for L436P, A398P+L436P, T435P and A398P+T435P mutants of Tau-5* were established using assignments from Tau-5* and allTau-5 CtoS PP.

### Binding studies

To evaluate EPI-001 binding to protein constructs, CSPs were determined by combining ^1^H and ^15^N chemical shift changes:

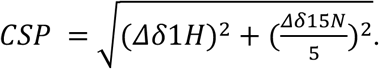

For NR ADs at 25 µM, we used the 2D ^1^H,^15^N FHSQC pulse sequence with WATERGATE as a water suppression module^64^ to assess CSPs in the absence and presence of 250 µM of EPI-001.

For other constructs we employed 2D ^1^H,^15^N BEST-TROSY. AR AD fragments (Nt, Tau-5*, Ct, and allTau-5 24G) and its mutant AR AD* (Extended Data Fig. 3e) were studied at a concentration of 25 µM. Tau-5* mutants were studied at 250 µM. Experiments were performed both in the absence and presence of EPI-001 (10 molar equivalents for AR AD fragments and 1 molar equivalent for Tau-5* mutants).

CSPs in the 2D ^1^H,^15^N BEST-TROSY spectra of 1Tau-5* and 2Tau-5* at a protein concentration of 0.31 mg/mL (25 µM 1Tau-5* and 12.5 µM 2Tau-5*) were measured in the presence of 250 µM EPI-001. For Tau-5* C404, CSPs induced by covalent binding with EPI-001 were examined at a protein concentration of 12 µM.

^1^H chemical shift changes of EPI-001 in the presence of the protein constructs were observed through 1D ^1^H experiments employing WATERGATE as the water suppression block^65^. Three to four distinct peaks of EPI-001 were monitored—two aromatic, a methylene, and a methyl group.

### MR AD competition with AR AD for EPI-001 binding

To assess competition between equimolar amounts of MR AD and AR AD for EPI-001 binding, two 2D ^1^H, ^15^N BEST-TROSY spectra of ^15^N-labelled AR AD (25 µM) plus 250 µM EPI-001 (10- fold molar excess) were compared: one in the absence and the other in the presence of ^14^N MR AD. A potential interaction between both proteins was ruled out by recording an additional 2D ^1^H, ^15^N BEST-TROSY spectra of AR AD (25 µM) in the presence of 1 molar equivalent of ^14^N MR AD.

### Helical content measurement

The helical content for AR AD and its fragments (Nt, Tau-5*, Ct, Extended Data Fig. 2f) was quantified at 25 µM protein, and at the assignment concentrations for the ADs of PR (200 µM), MR (200 µM), GR (280 µM), and ERɑ (150µM). For allTau-5 24G and Tau-5* constructs (WT, noAro, C404Y, A398P, G407A, and AAA), the concentration was set at 150 µM, unless otherwise specified in the figures. Tau-5* and Tau-5*:EPI-001 adduct were measured at 12 µM, with Tau- 5* featuring a single cysteine at position C404 in both cases (Tau-5* C404). For 1Tau-5* and 2Tau-5* it was obtained at 0.31 mg/mL.

^1^H, ^15^N, C’, and C_α_ chemical shifts obtained from 2D ^1^H, ^15^N correlation spectra and 3D HNCO and HNCA experiments were used as an input for the *δ*2D algorithm^51^. A 5% error was considered, as denoted in the red line of the helical content plots.

### CSP matrix

Unlabelled (^14^N) and isotopically labelled (^15^N) forms of protein constructs were mixed at a 200 µM concentration each. These experiments were carried out in the presence or absence of 200 µM EPI-001, with all samples containing 0.5% DMSO-d_6_, including all the samples in the absence of the small molecule. CSPs induced by adding 1 molar equivalent of unlabelled fragments were measured in 2D ^1^H,^15^N BEST-TROSY spectra of the labelled constructs (^15^N-Nt, ^15^N-Tau-5*, ^15^N-Ct, etc.). Detected CSPs reported on intermolecular interactions between fragments (e.g., ^15^N-Nt regions interacting with ^14^N-Nt, ^14^N-Tau-5*, or ^14^N-Ct).

We measured for each pair of interacting protein constructs the reciprocal experiments, i.e. the binding observed (NMR active nuclei, ^15^N) from each different construct. For a set of reciprocal datasets, e.g. ^15^N-Nt+^14^N-Tau-5* and ^15^N-Tau-5*+^14^N-Nt, we multiplied the two strings of CSP values to generate a m·n matrix, where m and n denote the length of the protein construct (e.g. for Nt+Tau5*, m_Nt_=151 and n_Tau5*_=119). The result of the product of two sets of CSP lists was defined as the contact parameter (*σ*). The value of *σ* for a residue i will increase if in the reciprocal dataset its CSP and the one corresponding to residue j (*σ*_ij_) are big. In IDPs, the dynamic nature facilitates promiscuous contacts but to ensure that certain interactions are taking place, mutations in specific regions were introduced to validate the contacts calculated. Also, a direct measurement of PRE NMR experiments, dependent on intermolecular distances, was conducted. The CSP matrix was represented as a heatmap, and a Gaussian filter was applied to average out missing data and emphasize regions with the highest CSPs caused by interactions.

Separate CSP matrices were computed for constructs both in the presence and absence of EPI-001 (or mutated helical motifs). Subsequently, the difference between these matrices was calculated (𝛥𝜎=*σ*_EPI-001_ - *σ*_apo_ or 𝛥𝜎=*σ*_no helix_ - *σ*_helix_). It’s important to note that if any of the corresponding values in one of the matrices (apo or with EPI-001) was missing or zero, the resulting position in the final matrix was set to zero. All data analysis was conducted with Python.

To confirm the changes in the CSP matrices (Extended Data Fig. 3c,d) are caused by disruptions in potential helical structure formation and not due to altered hydrophobicity resulting from the L436P mutation, we also evaluated the effect of the T435P mutation on the oligomerization of Tau-5*, which additionally contains the A398P mutation (Supplementary Information Fig. S3a).

### Paramagnetic relaxation enhancement

Intramolecular and intermolecular PREs for AR AD C404 were assessed by measuring 2D ^1^H,^15^N BEST-TROSY spectra of paramagnetic (para) and diamagnetic (dia) samples. The intensity ratio *I*_para_/*I*_dia_ was then calculated to quantify PREs.

For intramolecular PRE measurements, a sample containing 20 µM ^15^N-labelled AR AD with a paramagnetic spin label (MTSL) at C404 was mixed with 80 µM AR AD, where C404 was covalently blocked with iodoacetamide. Using this sample composition potential interactions between spin-labelled ^15^N AR AD and ^14^N AR AD would not produce PREs, minimizing but not completely preventing detection of intermolecular PREs. To account for intermolecular contributions, intermolecular PREs were measured at the same total protein concentration using a sample containing 20 µM ^15^N-labelled AR AD with C404 blocked with iodoacetamide, 20 µM AR AD spin-labelled at C404, and 60 µM AR AD with C404 blocked with iodoacetamide.

The diamagnetic state was achieved by adding 1.5 mM ascorbic acid for 24 hours at 4°C to reduce the paramagnetic spin label for both allTau-5 and AR C404S samples.

Intermolecular contacts of allTau-5 were measured on equimolar (100 µM) mixtures of spin-labelled ^14^N Cys-mutant allTau-5 (C404, C448 or C518) and ^15^N-labelled allTau-5 CtoS. 200 µM of EPI-001 was added to monitor the effects of the small molecule. The ^1^H^N^ *R*_2_ relaxation measurements were performed at 800 MHz at 5 °C. A total of six *T*_2_ relaxation times were used (1, 2, 5, 10, 20 and 60 ms). The ^1^H^N^ PREs were quantified by fitting the decay of signals to a single exponential function to obtain *R*_2*para,i*_ and *R*_2*dia,i*_ rates, from which the PRE contribution was calculated as *Γ*_2,*i*_ = *R*_2*para,i*_ − *R*_2*dia,i*_. Exponential curve fits were performed by using in-house-written scripts in R. Uncertainties in *R*_2_ were computed as the standard errors of the fit.

To visualise the strength of the measured non-covalent intermolecular interactions, we plotted the *Γ*_2_ values in the absence of the small molecule using the chord plot, as proposed previously^52^. In this data visualisation, each protein region is depicted as an arc (h2, ^391^LDYSAWAAAAAQ^403^; ^433^WHTLF^437^; C-term, C-terminal aromatic-rich (region 479-558) and the linkers connecting them) and the chords represent spatial proximity with a given spin-labelled position from the protein not enriched in ^15^N, depicted as red arcs. The thickness of the chords are proportional to the averaged *Γ*_2_ values (absence of EPI-001) or the Δ*Γ*_2_ (*Γ*_2,EPI-001_ - *Γ*_2,apo_) (to monitor the effect of EPI-001). The chord-plot representation provides quantitative information about the patterning of intermolecular contacts. The residue plots are presented in the Extended Data Fig. 3h.

Additionally, we analyzed intermolecular PREs of allTau-5 by plotting *I*_para_/*I*_dia_ ratios in the presence and absence of EPI-001, using spectra acquired with the 1 ms relaxation delay (Extended Data Fig. 3g). Standard errors for *I*_para_/*I*_dia_ values were estimated by propagating the intensity ratios of the individual spectra.

### Oligomerization studies

Oligomerization of 2Tau-5* and 1Tau-5* was analyzed by measuring CSPs in 2D ^1^H,^15^N BEST-TROSY experiments at a concentration of 0.31 mg/mL (25 µM 1Tau-5* and 12.5 µM 2Tau-5*). The samples also contained 0.5% DMSO-d_6_ to ensure consistency, as the same spectra were used for the EPI-001 binding studies.

To investigate the oligomerization of Tau-5* and its mutants, designed to modify aromaticity (noAro, h2SA, h3A, and C404Y) and helicity (A398P, L436P, G407A, and AAA), CSPs were monitored in 2D ^1^H,^15^N BEST-TROSY experiments of 25 µM and increasing the protein concentration up to 400 µM. The constructs employed in this analysis were either ^15^N labelled or ^15^N,^13^C-double labelled.

To confirm that the changes in Tau-5* oligomerization caused by the L436P mutation result from disruptions in potential helical structure formation rather than altered hydrophobicity, we also investigated the effect of the T435P mutation on Tau-5* oligomerization (Supplementary Information Fig. S3b).

### Cysteine *pK*_a_ measurements

The *pK*_a_ values of individual cysteine residues (C123, C404, and C518) were determined using ^15^N,^13^C-labelled single cysteine fragments: Nt C123, Tau-5* C404, and Ct C518. The cysteine ^13^C_β_ chemical shifts were measured in 70 µM Nt C123, 300 µM Tau-5* C404, and 150 µM Ct C518 samples. These measurements were conducted using ^1^H,^13^C-HSQC centered in the aliphatic region at varying *pH* from 6 to 12. All samples were prepared in the final buffer, containing 3 mM TCEP, to prevent disulfide bridge formation at high *pH* values.

The chemical shifts *δ*^13^C_β_ were fitted to the following equation, as previously reported^66^:

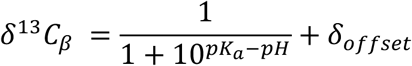

The *δ*^13^C_β_ scale was normalised (0<*δ*^13^C_β_<1) to correct for the slight differences in the chemical shift of the protonated state. The 95% confidence interval of the fitting was calculated by Monte Carlo error analysis.

### Dynamic Light Scattering

DLS measurements were performed using a Zetasizer Nano-S instrument (Malvern) equipped with a 633 nm He-Ne laser. All samples were freshly prepared prior to measurements, derived from stock solutions that had been filtered or centrifuged at 15,000 rpm for 10 minutes at 4 °C (the supernatant was used after the concentration was determined) and equilibrated for 10 minutes. Additionally, all samples of NR ADs and Tau-5* (except 1Tau-5* and 2Tau-5*) contained 0.5% DMSO-d_6_. Each sample was measured three times, and experiments were conducted at 5 °C. The percentage population of each species was primarily analyzed, as some samples exhibited a high polydispersity index (*pI*). The hydrodynamic radius *R*_h_ of monomers was confirmed by comparing experimental values to the theoretically calculated *R*_h_, as reported previously^67^.

### MicroScale Thermophoresis (MST)

Purified 2Tau-5* in a TCEP-free final buffer was labelled using the Monolith Protein Labeling Kit RED-NHS 2nd Generation (Nanotemper), following the manufacturer protocol. Measurements were performed on a Monolith NT.115 instrument, with data acquisition and analysis carried out using MO.Control and MO.Affinity Analysis softwares.

Labelled 2Tau-5* was used as the target at a concentration of 80 nM, with varying concentrations of EPI-001 (up to a solubility limit of 500 µM) at 25 °C. Samples contained 0.0125% Tween 20 and were loaded into Monolith Premium Capillaries (Nanotemper). The measurements were conducted at High MST power and 30% excitation power. Three measurements were performed for each sample, using independently prepared samples.

The binding curve was fitted to the analytical Hill equation to calculate the *K*_d_, assuming that the *F*_norm_ value at 500 µM represents the starting point of the bound plateau:

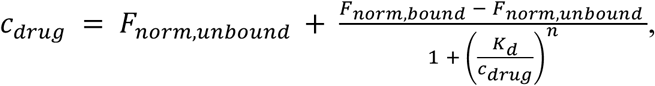

where *c*_drug_ represents the EPI-001 concentration, and *n* is the Hill coefficient.

### Mass spectrometry

25 µM protein samples were mixed with a 10-fold molar excess of the small molecule in the final buffer, and adjusted to *pH* 8. The mixtures were incubated at 37 °C from 1 to 24 hours. After the reaction, the *pH* of the samples was reduced to *pH* 7.4 using HCl. Subsequently, the samples were frozen and stored at -20 °C prior to their measurement by intact MS.

In Fig. 4e,f, single cysteine constructs were used —Tau-5* C404, Nt C123, Ct C518 and allTau-5 Cys mutants (C404, C448 and C518). For Fig. 4h, Tau-5* contained two cysteines - C404 and C448 - but no evidence of simultaneous covalent modification of both cysteines was observed. Another experiment using Tau-5* with a sole cysteine at C448 showed no covalent modification, indicating no covalent reaction takes place when the cysteine is at the C-terminus.

Frozen samples were thawed and diluted to a final concentration of 5 µM using a 3% v/v acetonitrile (ACN) and 1% v/v formic acid (FA) aqueous solution. Samples were analysed using an Acquity Ultra Performance chromatographic system coupled to an LCT Premier XE (TOF) mass spectrometer (Waters Corp., Milford, MA, US). 40 pmols of the sample were injected using the waters sample manager equipped with a Binary Solvent Manager. Protein content was separated on a BioSuite Phenyl 1000 column (RPC 2.0 × 75 mm, 10 µm, Waters Corp) with a linear gradient of 5% to 80% B (A= 0.1% v/v FA in water, B= 0.1% v/v FA in ACN). The column outlet interfaced directly with the electrospray ionisation (ESI) source of the spectrometer. The mass spectrometer operated in voltage analyzer mode with positive polarity. Capillary voltage and cone voltage were set at 3000 V and 100 V, respectively. Desolvation temperature and source temperature were 300 °C and 120 °C. Cone gas flow and desolvation gas flow were set at 50 and 600 l/h, respectively. Ion guide 1 and aperture 1 were set to 15 V and 10 V, respectively. Full MS scans (400 - 4000 m/z) were acquired using MassLynx software, V4.1.SCN704 (Waters Inc.). Manual deconvolution was performed using the MaxEnt 1 deconvolution algorithm, facilitated by V4.2.SCN982.

### Turbidity measurements

Turbidity measurements were performed to estimate *T*_c_ changes in allTau-5 induced by 250 µM EPI-001 or 5% allTau-5:EPI-001 adduct at a total protein concentration of 3 mg/mL and 100 mM NaCl.

To measure protein constructs *T*_c_, 2.4 mg/mL concentrations were used for Tau-5* mutants designed to modulate aromaticity and helicity, while maintaining 3 mg/mL for NR ADs and AR AD fragments, unless otherwise specified. All samples were pre-treated with 500 mM NaCl before experimentation. *T*_c_ values of AR AD* were used from the previous work^23^. Notably, we did not observe GR AD condensate formation even when the NaCl concentration was increased up to 2M. For Tau-5* mutants impacting aromatic content and patterning, we employed 1.5 M NaCl, needed to induce condensate formation of the h2SA and h3A mutants. The AR fragment Ct contained a modified shorter polyG sequence comprising only four glycines (Ct 4G) to prevent protein aggregation^58^.

The absorbances of the samples were recorded at 340 nm using 1 cm pathlength cuvettes and a Cary100 UV-Vis spectrometer equipped with a multicell thermoelectric temperature controller. A gradual temperature increase at a ramp rate of 1 °C/min was applied, and *T*_c_ were determined by identifying the peaks of the first-order derivatives of the curves. Standard deviations derived from three replicates were used to assess uncertainties.

### High performance liquid chromatography

The concentrations of the allTau-5:EPI-001 adduct and noAro, along with the *c*_sat_ measurement for allTau-5 and the determination of the partition coefficient of EPI-001 into protein condensates, were assessed using the Agilent Technologies 1260 Infinity II HPLC with a Jupiter analytical C4 column from Phenomenex. H_2_O and ACN:H_2_O (9:1) were used as mobile phases, containing 0.1 % v/v TFA.

### Saturation concentration measurement

Saturation concentration measurements were conducted to evaluate the alterations in the *c*_sat_ of allTau-5 induced by 250 µM EPI-001 or 100% allTau-5:EPI-001 adduct at a total protein concentration of 3 mg/mL. Formation of condensates was induced by 100 mM NaCl and samples were incubated at 25 °C for 5 minutes, followed by centrifugation for 2 minutes at 376 g. The supernatant was injected into the HPLC system, and the corresponding peaks representing allTau-5 or allTau-5:EPI-001 adducts at 49.2% ACN:H_2_O were integrated. Concentration was determined using calibration curves established for allTau-5 and allTau-5:EPI-001. The curves were generated by measuring the concentrations of individual samples using NanoDrop (Thermo Fisher Scientific) for allTau-5 and amino acid analysis for allTau-5:EPI-001. Subsequently, the samples and their dilutions were injected into the HPLC system to measure the peak integral dependency on concentrations.

### Measurement of small-molecule partition coefficients

To determine the partition coefficient of EPI-001 into the condensates of the constructs, samples with a protein concentration of 3 mg/mL were employed. For Tau-5* mutants influencing helicity (A398P, L436P, WT, G407A, and AAA), 250 µM EPI-001 was added, inducing condensation with 500 mM NaCl. In contrast, other constructs—Tau-5* mutants altering protein aromaticity (h2SA, h3A, WT, C404Y), AR AD fragments (AR AD, allTau-5, Tau-5*, and Ct 4G), and NR ADs (AR, MR, PR, ERɑ)—were supplemented with 100 µM EPI-001, and condensation was induced by 1.75 M NaCl. All samples were incubated for 5 minutes at 25 °C, followed by centrifugation 376 g for 2 min, allowing the separation of the light and dense phases. The light phase was transferred to a new microcentrifuge tube, while the dense phase was diluted with the final buffer containing 4 M urea to dissolve the condensates. Both dense and light phases were injected into the HPLC system, and the EPI-001 signal was detected at 49.4% of ACN:H_2_O. Integrating the corresponding peak of the ligand enabled the calculation of the ratio between the dense and light phases, determining the partition coefficient. Notably, partition coefficient measurements were not conducted for the Nt fragment and GR AD due to the absence of observed condensate formation at concentrations up to 2 M NaCl.

### LacI tethering assay

The activation domains of ERɑ, MR, and GR constructs were ordered from Twist Bioscience. The AD of AR and FUS LCD were amplified from plasmids utilised in our previous work^23^, created in the D. Hnisz lab (Addgene: 215637), respectively. The AD of PR was amplified from MCF7 cDNA. Primers used for the construct amplification are listed in the primer table in Table S2 (Supplementary Information). The amplified constructs were then inserted into the LacI-CFP-MCS plasmid pJM118^42^ using the NEBuilder HiFi DNA Assembly Master Mix kit (NEB E2621X). The insert:vector ratio used for assembly was 1:3 with 45-50 ng of vector. Cloned constructs were validated using Sanger sequencing.

25,000 U2OS cells with a lacO array integrated into the genome^39^ were seeded per well in Ibidi 8-well glass bottom u-slides. The following day, cells were transfected with 100 ng of YFP-containing plasmid (YFP empty for control or YFP-RNAP II CTD (Naderi et al., NCB, 2024, accepted)) and 75 ng of lacI-CFP(AD of NR or FUS LCD) plasmid using Lipofectamine 3000 (ThermoFischer Scientific) according to the manufacturer protocol. 24 h after transfection, the medium was changed to medium containing either DMSO or 25 µM EPI-001. Cells were imaged approximately 24 h after DMSO or EPI-001 treatment using a Zeiss LSM880 confocal microscope with a live cell imaging chamber set to 37 °C and 5% CO_2_, using a Plan-Apochromat-63x/1.40 oil DIC objective with a 2x zoom. Two biological replicates with 18-22 images were taken for each condition per replicate.

Acquired images were analysed using FIJI^68^. YFP and CFP intensities were measured in CFP tethers, which were manually selected for each cell. An area of the same size and shape was utilised for YFP background level normalisation and enrichment calculation (intensity in tether/intensity in background) in each cell. The data presented in Fig. 6d was replotted for Fig. 6e, specifically for the DMSO with RNAP II CTD condition.

### Fluorescence microscopy

Purified proteins, including AR AD, allTau-5 C518, and Tau-5* C404, were labelled with DyLight 488 dye featuring Maleimide thiol (Thermo Fisher Scientific). The protein labelling process adhered to the manufacturer protocol. The proteins and the dye were mixed at an approximate ratio of 1:10 in the final protein solution and left overnight at 4°C. Subsequently, isolation of the protein and labelled protein from the unbound dye was accomplished through SEC using HiLoad superdex 75 or 200 pg 13/300 columns. Concentrations and conjugation efficiencies were assessed following the manufacturer guidelines. For fluorescence microscopy samples, a 1 µM dye concentration was employed.

For fluorescence microscopy imaging, AR AD fragments (AR AD, allTau5, and Tau-5*) were prepared at a concentration of 3 mg/mL, while Tau-5* mutants (C404Y, G407A, and WT) were at 2.4 mg/mL (200 µM). These proteins were imaged both in the presence and absence of 250 and 200 µM of EPI-001, respectively. Condensate formation was induced by adding 500 mM NaCl. In a chamber made of a slide and a coverslip with double-sided tape (3M 300 LSE high-temperature double-sided tape, 0.17 mm thickness), 1.8 µL of the sample was deposited. The coverslips were coated with PEG-silane as per the protocol by Alberti et al. (2018). Imaging was performed on the coverslip surface where the droplets were settled using a Zeiss LSM780 Confocal Microscope equipped with a Plan-ApoChromat x63/1.4 Oil objective lens. Laser power was adjusted to optimize the visualization of droplets. Experiments involving AR AD fragments were conducted at 37 °C, while those involving Tau-5* mutants were performed at 30 °C.

Equivalent-sized droplets were selected for fluorescence recovery after photobleaching (FRAP) analysis, with the bleached area covering approximately 30% of their diameter. Intensity values were monitored across different regions of interest (ROIs): ROI1 corresponded to the bleached area, ROI2 encompassed the entire droplet, while ROI3 captured the background signal. The obtained data underwent processing via EasyFrap software, enabling the extraction of kinetic parameters such as the half-time of recovery (*t*_1/2_)^69^.

### Statistical analysis

Pairwise comparisons were conducted using the t-test, except for the data in Fig. 5d,e and Extended Data Extended Data Fig. 5c,d, for which Wilcoxon tests were used for statistical analysis. Significance levels were determined based on the p-values generated by the test, denoting significance as follows: **** (*p* < 0.0001), *** (*p* < 0.001), ** (*p* < 0.01), or * (*p* < 0.05).

## Acknowledgments

We thank all members of the Laboratory of Molecular Biophysics at IRB, Nuage Therapeutics, and the Multi-Level Gene Control Lab at the Max Planck Institute for Molecular Genetics for their insightful discussions. We thank I. Latorre, J. M. Valverde and J. Lluna for their assistance with protein production. We thank L. Pont Villanueva and F. J. Benavente Moreno from Barcelona University, and to P. Pheifer and G. Battaglia from IBEC, for their introduction to and assistance with the HPLC experiments. We thank the support and assistance provided by the Advanced Digital Microscopy and Mass Spectrometry facilities at IRB Barcelona, particularly M. del Mar Vilanova. We thank the support received from the Spanish ICTS Network of Biomolecular NMR Laboratories (R-LRB).

S.B. acknowledges support from a PhD fellowship awarded by IRB in the 2020 call and an FPI fellowship awarded by MINECO in the 2021 call (PID2019-110198RV-100). L.B. is supported by PRIN 2022 (2022EMZJL4). B.M. is supported by an AECC (Spanish Cancer Association) postdoctoral grant. D.H. acknowledges support by the Mark Foundation for Cancer Research through an ASPIRE award. X.S. was supported by AGAUR (2021 SGR 476), MICINN (PID2019-110198RB-I00), the Fundació La Caixa (CI20-00098), the AECC (INNO20010FRIG) and the European Research Council (CONCERT, contract number 648201). A.R. was supported by MICINN (PID2020-115074GB-I00). IRB Barcelona is the recipient of a Severo Ochoa Award of Excellence from MINECO (Government of Spain).

## Author contributions

Conceptualization: S.B., B.M. and X.S.; investigation: S.B., B.M., M.A., H.N., C.S.Z., C.G.C., L.B., J.G.; visualisation: S.B. and B.M.; methodology: S.B., B.M., M.A., H.N., M.F.V., J.G.; project administration: S.B. and B.M.; supervision: X.S. with contribution from R.P., I.C.F., A.R. and D.H.; funding acquisition: A.R., D.H. and X.S.; writing – original draft: S.B., B.M. and X.S.; writing – review & editing: all authors.

## Supplementary information

Tables S1-S2

Fig. S1-S5

## Extended Data

**Extended Data Fig. 1.**
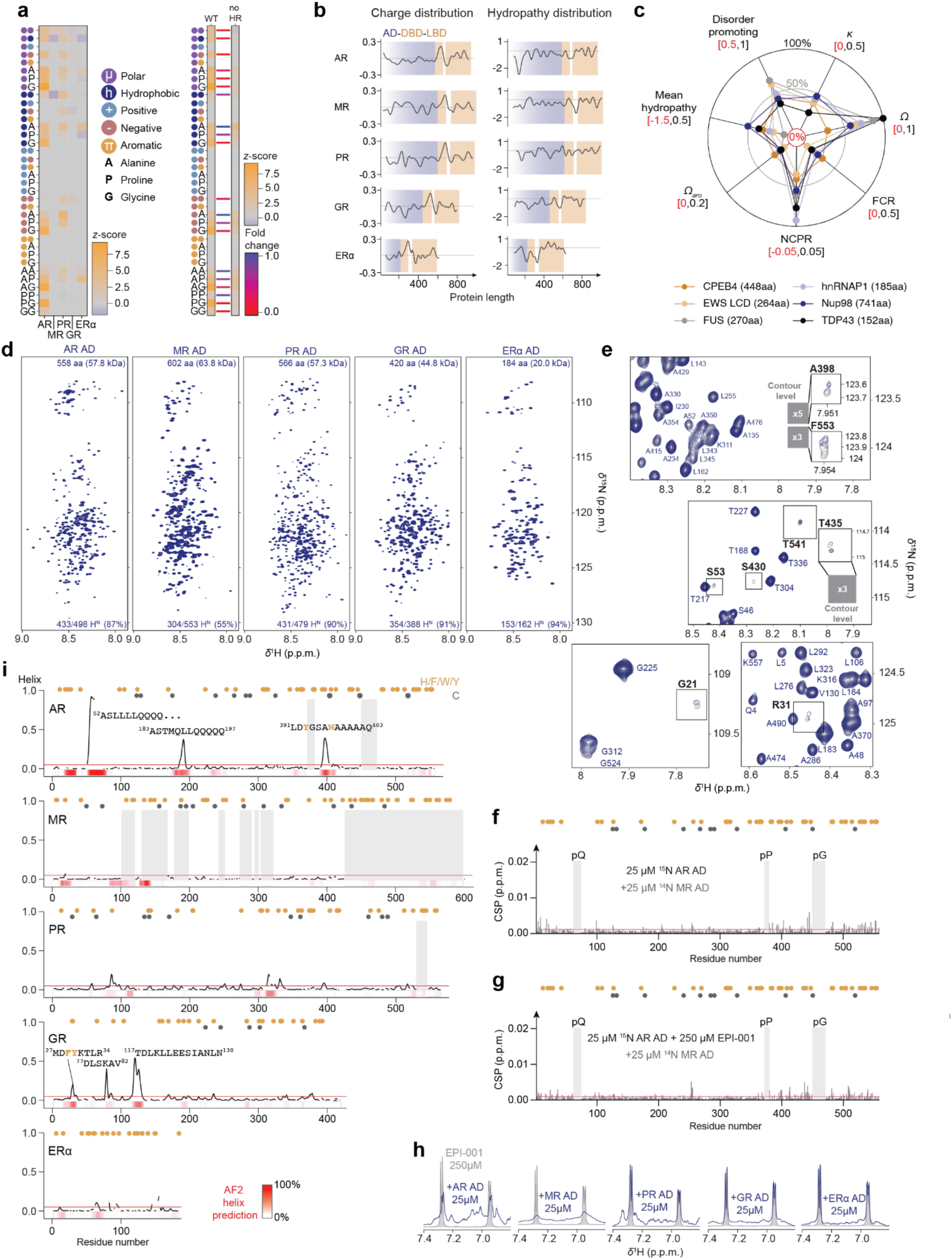
(**a**) (*Left*) Clustering analysis of non-random amino acid sequences across the activation domains of the nuclear receptor family^53^. Positive *z*-scores (orange) indicate a linear clustering (“blocky”), and negative values (blue) indicate uniform dispersion (“well-mixed”). These parameters were calculated for residue classes that are present in a proportion that is greater than 10%. (*Right*) NARDINI analysis^53^ of the sequence segregation of certain amino acid-type classes in AR AD in the presence and absence of homorepeats (poly-glutamine, -proline and -glycine). Positive *z*-scores (orange) indicate a blockiness of these amino acids compared to a random distribution of *in silico* generated 10^5^ sequences that retain the overall composition. In general, the sequence segregation is reduced by eliminating the homorepeats. (**b**) Sequence profile of the physicochemical properties (charge and hydropathy). (**c**) Overall sequence properties for the IDRs of TDP-43, FUS, EWS low-complexity domain, Nup98, hnRNAP1, and CPEB4. Mean hydropathy; Fraction of disorder promoting residues in the sequence; *κ* and *Ω* values describe the segregation of charged and proline residues^54,55^; FCR, fraction of charged residue; NCPR, net charge per residues; Aromatic clustering measures the normalized patterning of aromatic residues^56^. (**d**) 2D ^1^H-^15^N NMR correlation spectra of the different NR ADs. The total amount of amide protons assigned, either from sequential assignment or indirectly from a divide-and-conquer approach or urea titration, is indicated. The narrow dispersion in the ^1^H dimension is an indicative NMR signature of IDRs. The heterogeneous intensities are a typical property of IDRs mediating intermolecular homotypic interactions with other protein molecules. **(e)** Representative NMR signals of the AR AD (grey) that experience significant CSP in the presence of 10 molar equivalents of EPI-001 (blue) as shown in Figure 1e. (**f**) CSPs extracted from 2D ^1^H-^15^N NMR correlation spectra of AR AD induced by 1 molar equivalent of MR AD. (**g**) CSPs induced by MR AD in the 2D ^1^H-^15^N NMR correlation spectra of 1 molar equivalent of AR AD and 10 equimolar EPI-001. (**h**) Aromatic regions of 1D ^1^H spectra of EPI-001 in the absence (grey) or presence (blue) of a 0.1 molar equivalent of the corresponding AD. (**i**) NMR-derived helical content of NR ADs. The red line indicates the uncertainty of the *δ*2D algorithm^51^. Helical populations for AR AD were extracted from the chemical shift of separate constructs^23^. AlphaFold predicted helical populations are indicated as a red barcode in the bottom of each plot. (**f, g**) The red line represents the significant threshold calculated as the average plus five standard deviations of the first quartile of CSP. (**f, g, i**) Orange and grey dots above the residue plots indicate the position of aromatic or cysteine residues, respectively. Grey shadow squares represent the regions that have an incomplete assignment due to excessive line broadening or signal overlap.

**Extended Data Fig. 2.**
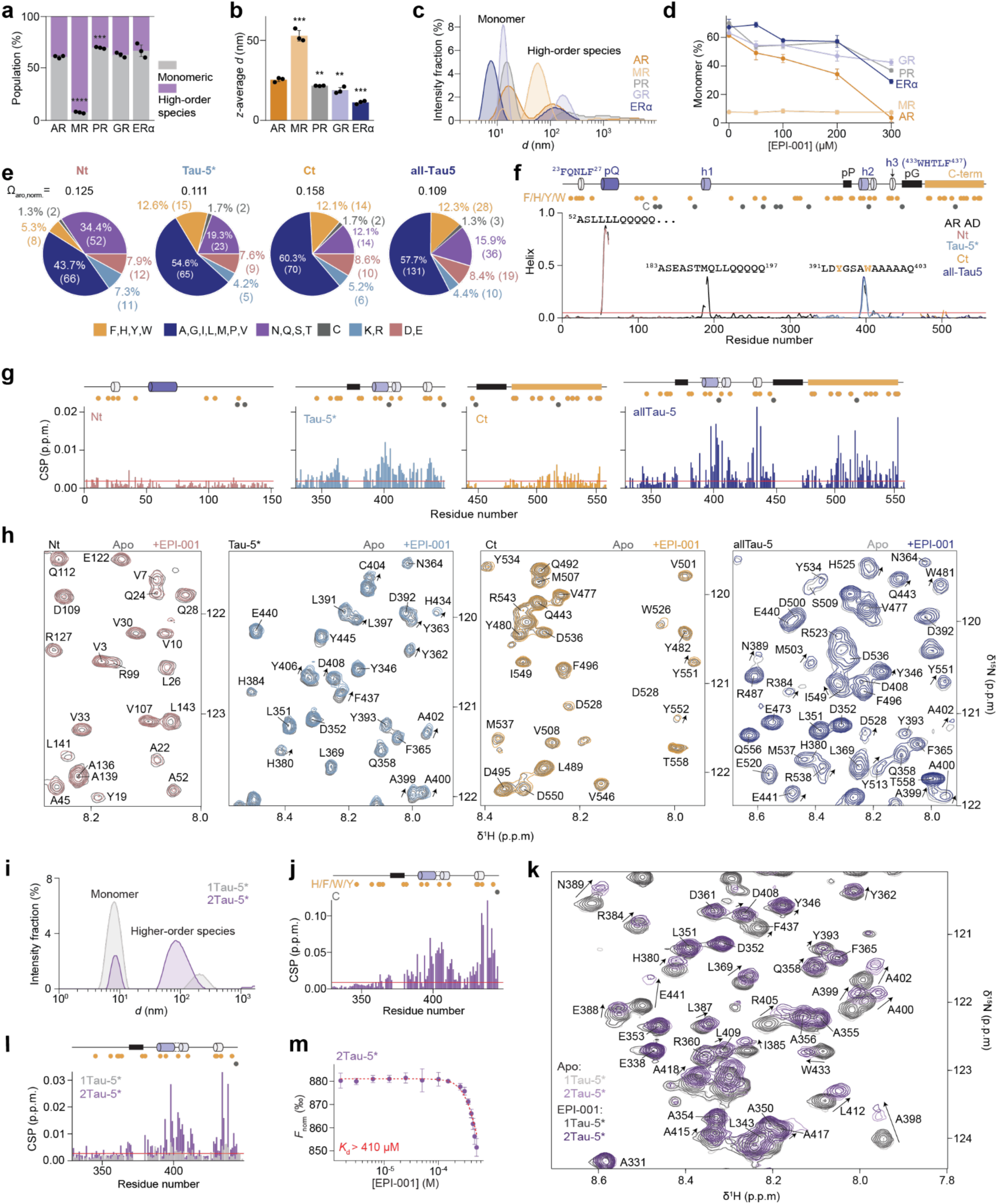
(**a**) Percentage of monomeric and higher-order species and (**b**) Z-average hydrodynamic diameter in NR AD samples analyzed by DLS at 2 mg/mL. (**c**) Corresponding DLS intensity measurement of NR ADs. (**d**) Percentage of monomer in 2 mg/mL NR AD samples at varying concentrations of EPI-001. The remaining percentage represents high-order species measured by DLS. (**e**) Amino acid composition of AR AD fragments depicted for five different residue types: polar (purple), cysteine (grey), positive charge (light blue), negative charge (pink), aromatic (orange), and hydrophobic (blue). (**f**) Annotation of short helical motifs in the AR AD. The plots display the helical propensity of the AR AD and its fragments, measured by NMR. The ^23^FQNLF^27^ and ^433^WHTLF^437^(h3) motifs are known to fold upon binding to cellular partners^28^. Orange and grey circles indicate the positions of aromatic and cysteine residues, respectively. The red line indicates the uncertainty of the algorithm (5%)^51^. (**g**) CSPs induced by 250 µM EPI-001 were measured in 2D ^1^H-^15^N NMR correlation spectra of AR AD fragments at a protein concentration of 25 µM. (**h**) NMR spectral region of the AR AD fragments in the presence (colour) and absence (gray) of the compound. (**i**) DLS intensity measurement of 1Tau-5* and 2Tau-5* at a concentration of 2.44 mg/mL. (**j**) CSPs obtained by comparing the 2D ^1^H-^15^N NMR correlation spectra of 1Tau-5* and 2Tau-5* at 0.31 mg/mL. (**k**) Representative NMR signals of 1Tau-5* and 2Tau-5* in the absence and presence of EPI-001. (**l**) Per-residue CSPs in 2D ^1^H-^15^N NMR correlation spectra of 1Tau-5* and 2Tau-5* at 0.31 mg/mL after adding 250 µM EPI-001. In both cases, the ratio corresponds to 1:10 EPI-001 interaction sites on the protein to EPI-001. (**m**) MST binding curves of EPI-001 with 2Tau-5* at a fixed protein concentration of 80 nM. The curve was fitted using the analytical Hill equation, and the *K*_d_ value was calculated under the assumption that a plateau is reached at the highest measured concentration of EPI-001. MST traces are presented in Supplementary Information: Fig. S5. (**a, b, d, m**) Error bars indicate the standard deviations (*n* = 3). (**g, j, l**) The red line represents the significant threshold calculated as the average plus five standard deviations of the first quartile of CSPs.

**Extended Data Fig. 3.**
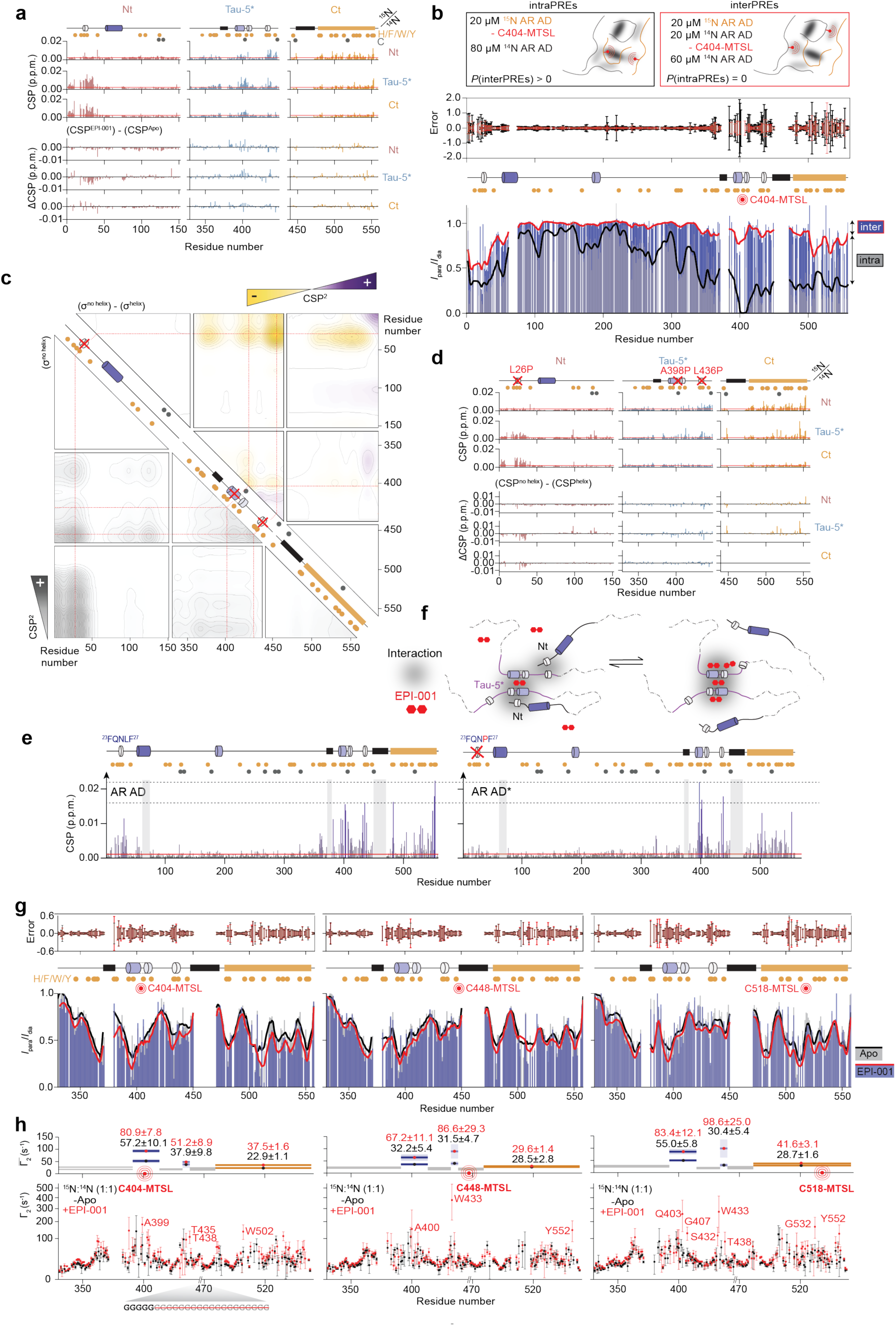
(**a**) (*Top*) CSPs residue plots obtained from 2D ^1^H-^15^N NMR correlation spectra, used to calculate the contact map in the absence of EPI-001 and (*bottom*) difference CSP residue plots used to calculate the Δ*σ* values shown in Fig. 3a.Orange and grey circles indicate the positions of aromatic and cysteine residues, respectively. (**b**) Intra and intermolecular interactions within AR AD analyzed using PRE NMR experiments. *(Top)* Schematic of the experimental design. "*P*" represents the probability of detecting inter-PREs arising from interactions between two ^15^N-labelled AR AD molecules in an experiment designed to measure intramolecular interactions, and intra-PREs in an experiment designed to detect intermolecular interactions. *(Middle)* Error bars are inversely proportional to the propagated signal-to-noise ratio of individual resonances. *(Bottom)* PRE intensity ratio graphs (*I*_para_/*I*_dia_) illustrate intermolecular and intramolecular proximity in AR AD with a paramagnetic spin label at the C404 position, while all other cysteines were replaced by serines. (**c**) CSP matrix between equimolar mixtures AR AD fragments where the helical elements were mutated (L26P in Nt, A398P and L436P in Tau-5*). CSP^2^ (*σ*) in the absence of helical content are shown in grey. Differences in the CSP^2^ (Δ*σ*) upon transient helix formation are shown in the upper matrix. (**d**) CSPs residue plots, obtained from 2D ^1^H-^15^N NMR correlation spectra, used to generate the CSP matrices shown in Extended Data Fig. 3c. (**e**) Per-residue CSPs induced by 10 molar equivalents of EPI-001 in AR AD and AR AD*^23^ 2D ^1^H-^15^N NMR correlation spectra. AR AD* contains the L26P mutation, which prevents secondary structure formation within the ^23^FQNLF^27^ motif. (**f**) Scheme illustrating EPI-001 competition with the ^23^FQNLF^27^ motif for binding to its target site. (**g, h**) Intermolecular interactions between allTau-5 molecules monitored via PRE NMR experiments in the absence (black) and presence of 1 molar equivalent of the ligand (red). (**g**) *(Top)* Error bars are inversely proportional to the propagated signal-to-noise ratio of individual resonances. *(Bottom)* PRE intensity ratio graphs (*I*_para_/*I*_dia_) illustrate intramolecular interactions in allTau-5 with a paramagnetic spin label at different cysteine positions, while remaining cysteines were mutated to serine. (**h**) Error bars indicate the standard error from the exponential fit. The average *Γ*_2_^HN^ for certain regions are shown in upper plots. Error bars represent the standard deviations. Differences in PREs between the presence and absence of EPI-001 are shown in Supplementary Information: Fig. S4. (**a, d, e**) The red line represents the significant threshold calculated as the average plus five standard deviations of the first quartile of CSPs.

**Extended Data Fig. 4.**
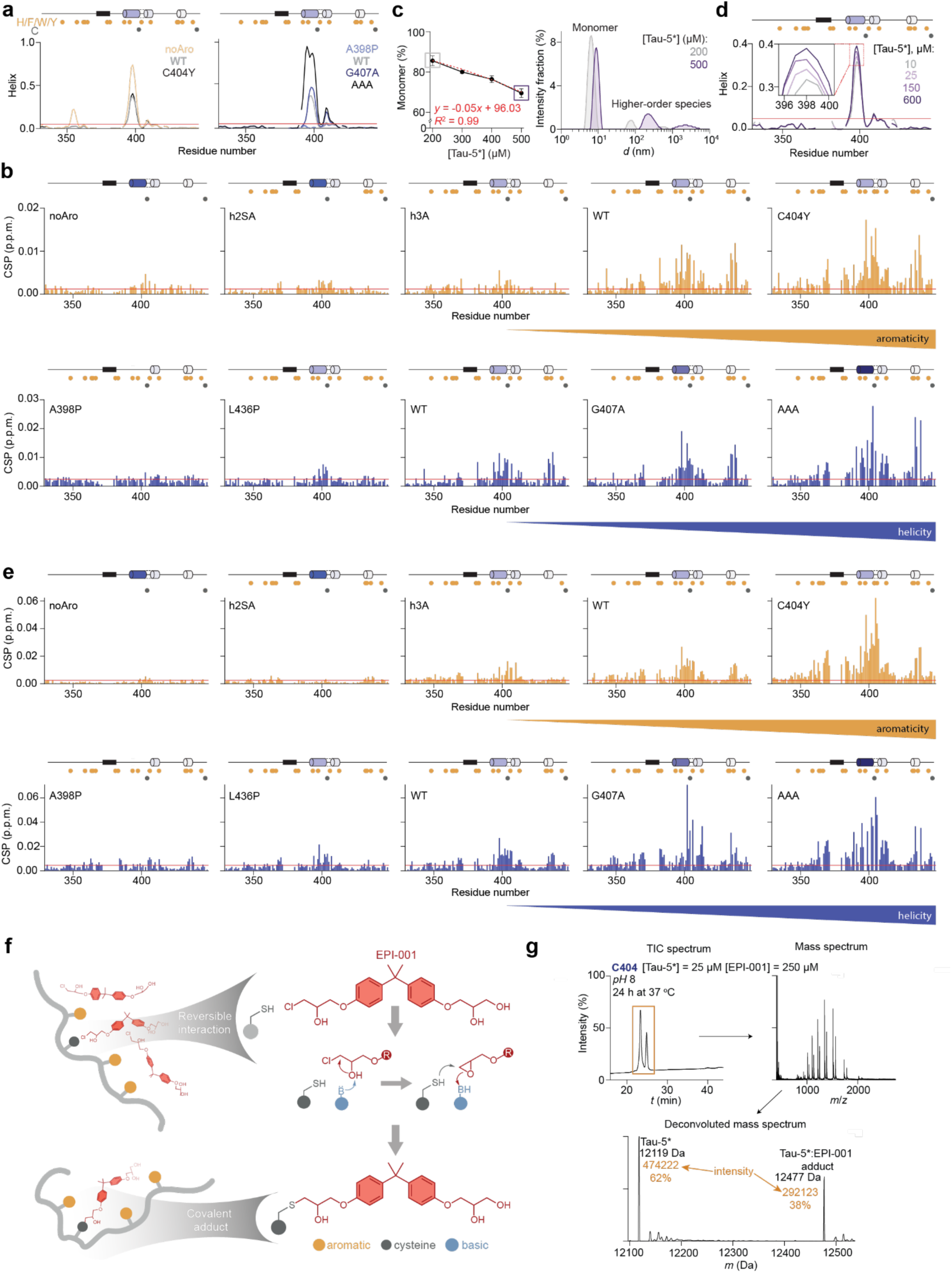
(**a**) NMR-derived helical content of Tau-5* mutants. Orange and grey dots indicate the positions of aromatic and cysteine residues, respectively. (**b**) CSPs detected in the 2D ^1^H-^15^N NMR correlation spectra of Tau-5* mutants, engineered to modulate aromaticity or helicity, in response to 1 molar equivalent of EPI-001. (**c**) (*Left*) Percentage of monomer at different concentrations of Tau-5* samples measured by DLS, with the remaining percentage corresponding to higher-order species. (*Right*) DLS intensity measurement of Tau-5* at concentrations of 200 µM and 500 µM. Error bars indicate the standard deviations (*n* = 3). (**d**) NMR-derived helical content of Tau-5* at various concentrations. (**e**) CSPs detected in 2D ^1^H-^15^N NMR correlation spectra of Tau-5* mutants induced by increasing concentrations of the protein from 25 to 400 µM. These CSPs indicate regions involved in protein oligomerization. (**f**) Scheme showing the covalent reaction mechanism between the chlorohydrin warhead and the thiol group of cysteines. In a first step, the proximity between both chemical groups is induced by the transient binding of the aromatic side-chains. (**g**) Schematic workflow of the quantification of covalent adduct formation by HPLC-MS. The adduct and the free protein were separated by liquid chromatography and the mass of each peak was determined by intact MS. The peak intensity (integral) of the deconvoluted spectra was used to calculate the relative populations of protein and adduct. (**a, d**) The red line indicates the uncertainty of the *δ*2D algorithm (5%)^51^. (**b, e**) The red line represents the significant threshold calculated as the average plus five standard deviations of the first quartile of CSPs.

**Extended Data Fig. 5.**
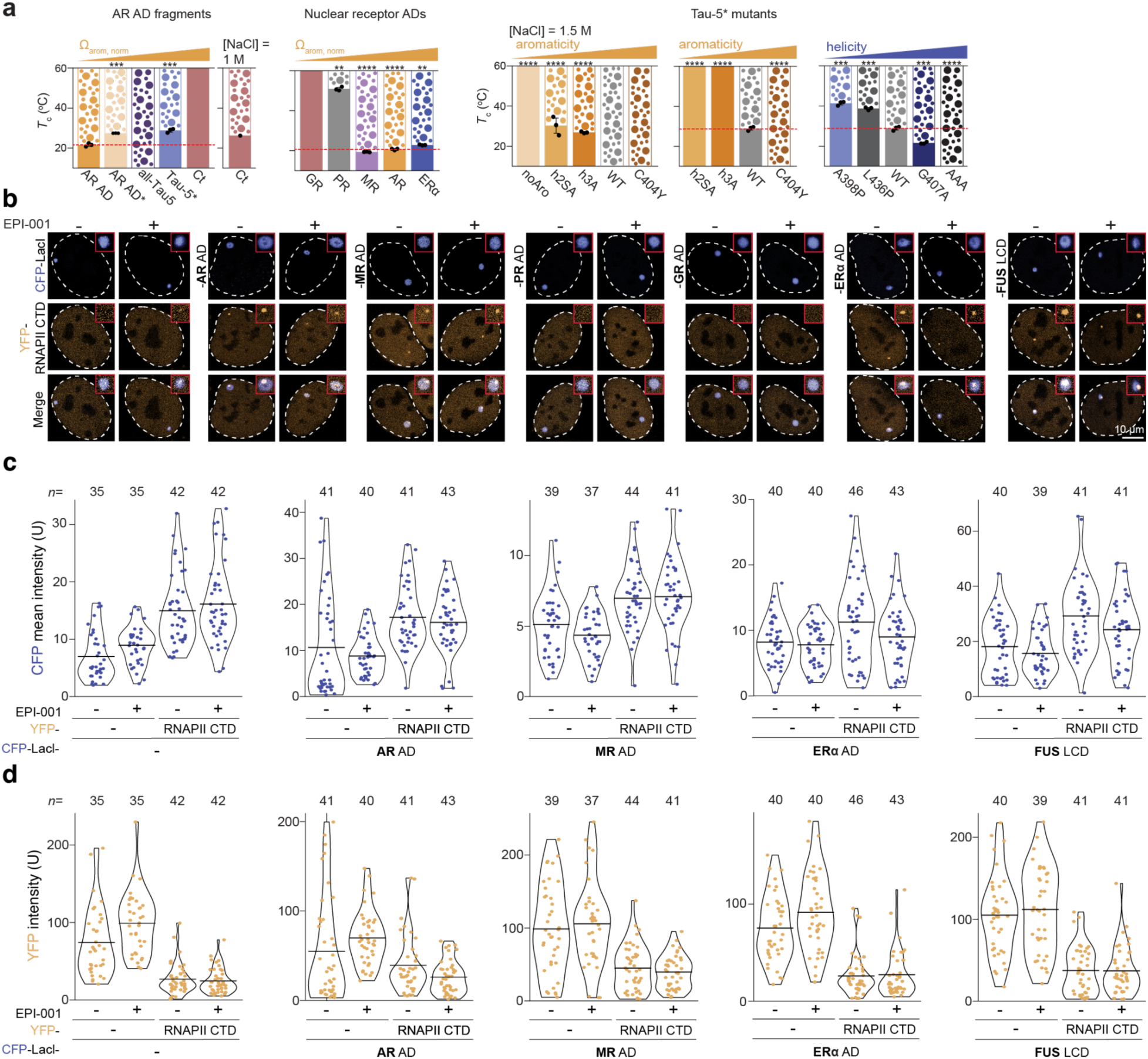
(**a**) *T*_c_ of AR AD fragments, NR ADs and Tau-5* mutants at 500 mM NaCl, unless otherwise specified. Errors shown on dots are standard deviations (*n* = 3). (**b**) Representative images of cells co-transfected with YFP-RNAP II CTD and CFP-LacI-(NR AD or FUS LCD). Cells were treated either in the absence(-) or presence(+) of 25 µM EPI-001. Each image depicts one example nucleus, with the nuclear contour highlighted by a white dashed line. Within the red squares, a zoomed-in version of the CFP focus is displayed. Notably, the RNAPII CTD signal does not entirely overlap with the tether, consistent with known characteristics of RNAPII CTD recruitment in this assay^40,41^. (**c**) Comparative analysis of CFP signal intensity within the tethers of the NR ADs and FUS LCD foci. (**d**) Comparison of YFP(-) and RNAP II CD-YFP signal intensities in the cellular background of cells transfected with different NR ADs and FUS LCD tethers. (**c, d**) Each data point represents a single tether/cell.

**Extended Data Fig. 6.**
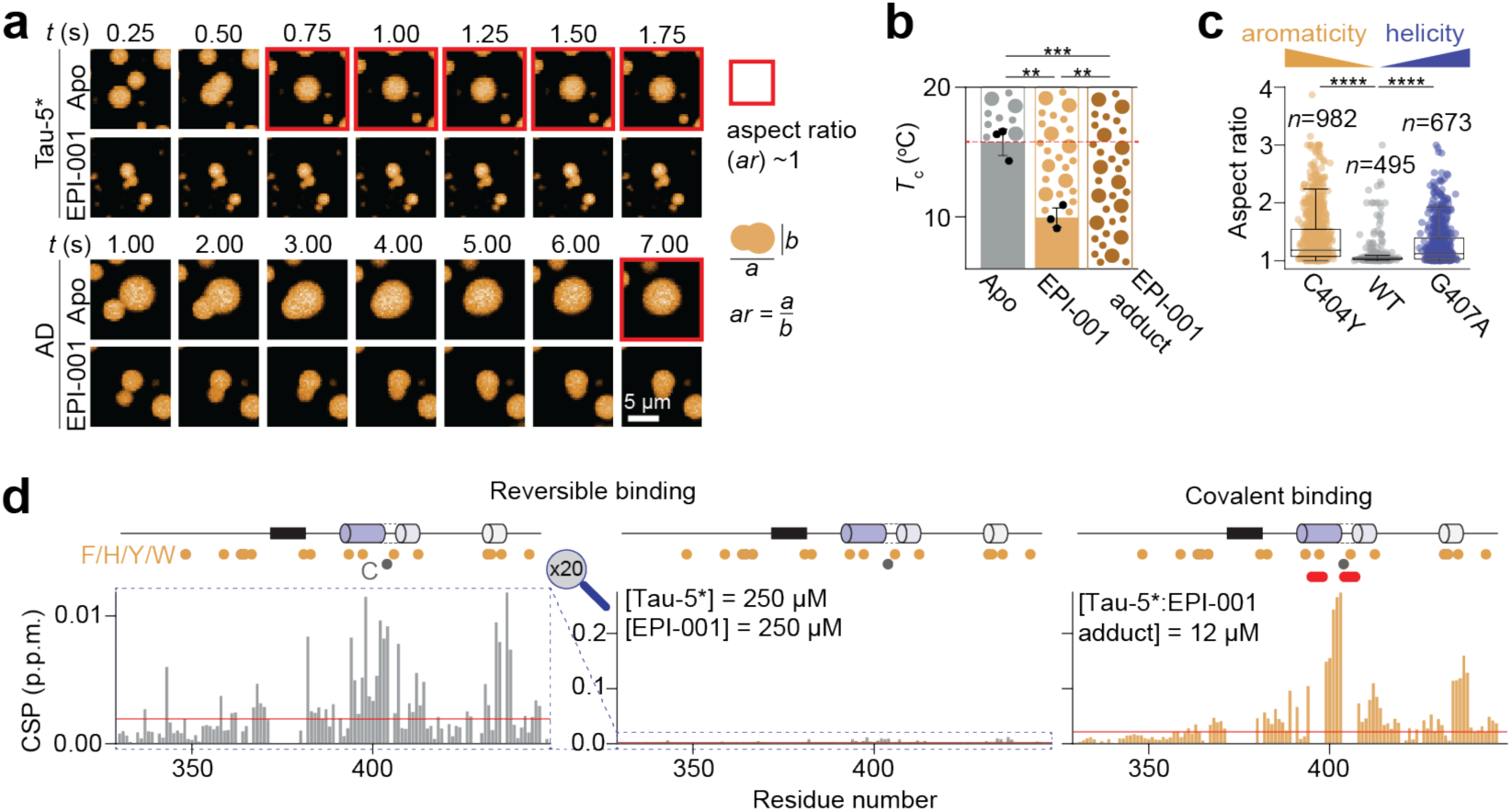
(**a**) Fusion events monitored at different time points of AR AD and Tau-5* condensates in the presence and absence of EPI-001 *in vitro*. Spherical fused droplets are highlighted in red. (**b**) *T*_c_ measurements of allTau-5 in the presence and absence of EPI-001 (ratio 1:1), and allTau-5*:EPI-001 adduct. Reduced *T*_c_ indicates an enhancement of condensation by the ligand. Errors shown on dots are standard deviations (*n* = 3). (**c**) CSPs in 2D ^1^H-^15^N NMR correlation spectra of Tau-5* induced by the covalent and reversible interaction of 1 molar equivalent EPI-001, respectively. Covalent binding induced CSPs approximately 20 times higher than reversible interaction. Orange and grey circles indicate the positions of aromatic and cysteine residues, respectively. Red circles and indicate not assignable residues due to line broadening. The red line represents the significant threshold calculated as the average plus five standard deviations of the first quartile of CSPs. (**d**) Aspect ratio quantification of Tau-5* mutants based on fluorescence microscopy images shown in Fig. 6g.

