## Supplementary information for "Oligomerization enables the selective targeting of intrinsically disordered regions by small molecules"

**Table S1.** The sequences of proteins used for the in vitro experiments. Mutation positions are in bold font and underlined.

| Name | Protein sequence |
|---|---|
| AR AD | $^{1}$MEVQLGLGRVYPRPPSKTYRGAFQNLFQSVREVIQNPGPRHPEAASAAPPGASLLLLQQQQQ<br>QQQQQQQQQQQQQQQQETSPRQQQQQQGEDGSPQAHRRGPTGYLVLDEEQQPSQPQSALE<br>CHPERGCVPEPGAAVAASKGLPQQLPAPPDEDDSAAPSTLSLLGPTFPGLSSCSADLKDILSEA<br>STMQLLQQQQQEAVSEGSSSGRAREASGAPTSSKDNYLGGTSTISDNAKELCKAVSVSMGLG<br>VEALEHLSPGEQLRGDCMYAPLLGVPPAVRPTPCAPLAECKGSLLDDSAGKSTEDTAEYSPF<br>KGGYTKGLEGESLGCSGSAAAGSSGTLELPSTLSLYKSGALDEAAAYQSRDYYNFPLALAGP<br>PPPPPPPHPHARIKLENPLDYGSAWAAAAAQCRYGDLASLHGAGAAGPGSGSPSAAASSSWH<br>TLFTAEEGQLYGPCGGGGGGGGGGGGGGGGGGGGGGGGEAGAVAPYGYTRPPQGLAGQES<br>DFTAPDVWYPGGMVSRVPYPSPTCVKSEMGPWMDSYSGPYGDMRLETARDHVLPIDYYFPP<br>QKT$^{558}$ |
| AR AD* (L26P) | $^{1}$MEVQLGLGRVYPRPPSKTYRGAFQN**P**FQSVREVIQNPGPRHPEAASAAPPGASLLLLQQQQQ<br>QQQQQQQQQQQQQQQQETSPRQQQQQQGEDGSPQAHRRGPTGYLVLDEEQQPSQPQSALE<br>CHPERGCVPEPGAAVAASKGLPQQLPAPPDEDDSAAPSTLSLLGPTFPGLSSCSADLKDILSEA<br>STMQLLQQQQQEAVSEGSSSGRAREASGAPTSSKDNYLGGTSTISDNAKELCKAVSVSMGLG<br>VEALEHLSPGEQLRGDCMYAPLLGVPPAVRPTPCAPLAECKGSLLDDSAGKSTEDTAEYSPF<br>KGGYTKGLEGESLGCSGSAAAGSSGTLELPSTLSLYKSGALDEAAAYQSRDYYNFPLALAGP<br>PPPPPPPHPHARIKLENPLDYGSAWAAAAAQCRYGDLASLHGAGAAGPGSGSPSAAASSSWH<br>TLFTAEEGQLYGPCGGGGGGGGGGGGGGGGGGGGGGGGEAGAVAPYGYTRPPQGLAGQES<br>DFTAPDVWYPGGMVSRVPYPSPTCVKSEMGPWMDSYSGPYGDMRLETARDHVLPIDYYFPP<br>QKT$^{558}$ |
| AR C404 | $^{1}$MEVQLGLGRVYPRPPSKTYRGAFQNLFQSVREVIQNPGPRHPEAASAAPPGASLLLLQQQQQ<br>QQQQQQQQQQQQQQQQETSPRQQQQQQGEDGSPQAHRRGPTGYLVLDEEQQPSQPQSALE*<br>*HPERG*VPEPGAAVAASKGLPQQLPAPPDEDDSAAPSTLSLLGPTFPGLSS*SADLKDILSEAS<br>TMQLLQQQQQEAVSEGSSSGRAREASGAPTSSKDNYLGGTSTISDNAKEL*KAVSVSMGLGV<br>EALEHLSPGEQLRGD*MYAPLLGVPPAVRPTP*APLAE*KGSLLDDSAGKSTEDTAEYSPFKG<br>GYTKGLEGESLG*SGSAAAGSSGTLELPSTLSLYKSGALDEAAAYQSRDYYNFPLALAGPPPP<br>PPPPHPHARIKLENPLDYGSAWAAAAAQCRYGDLASLHGAGAAGPGSGSPSAAASSSWHTLF<br>TAEEGQLYGP*GGGGGGGGGGGGGGGGGGGGGGGGEAGAVAPYGYTRPPQGLAGQESDFT<br>APDVWYPGGMVSRVPYPSPT*VKSEMGPWMDSYSGPYGDMRLETARDHVLPIDYYFPPQKT<br>$^{558}$ |
| MR AD | $^{1}$METKGYHSLPEGLDMERRWGQVSQAVERSSLGPTERTDENNYMEIVNVSCVSGAI<br>PNNSTQGSSKEKQELLPCLQQDNNRPGILTSDIKTELESKELSATVAESMGLYMDSV<br>RDADYSYEQQNQQGSMSPAKIYQNVEQLVKFYKGNGHRPSTLSCVNTPLRSFMSD<br>SGSSVNGGVMRAVVKSPIMCHEKSPSVCSPLNMTSSVCSPAGINSVSSTTASFGSFP<br>VHSPITQGTPLTCSPNVENRGSRSHSPAHASNVGSPLSSPLSSMKSSISSPPSHCSVKS<br>PVSSPNNVTLRSSVSSPANINNSRCSVSSPSNTNNRSTLSSPAASTVGSICSPVNNAFS<br>YTASGTSAGSSTLRDVVPSPDTQEKGAQEVPFPKTEEVESAISNGVTGQLNIVQYIK<br>PEPDGAFSSSCLGGNSKINSDSSFSVPIKQESTKHSCSGTSFKGNPTVNPFPFMDGSY<br>FSFMDDKDYYSLSGILGPPVPGFDGNCEGSGFPVGIKQEPDDGSYYPEASIPSSAIVG<br>VNSGGQSFHYRIGAQGTISLSRSARDQSFQHLSSFPPVNTLVESWKSHGDLSSRRSD<br>GYPVLEYIPENVSSSTLRSVSTGSSRPSKI$^{602}$ |
| PR AD | $^{1}$MTELKAKGPRAPHVAGGPPSPEVGSPLLCRPAAGPFPGSQTSDTLPEVSAIPISLDG<br>LLFPRPCQGQDPSDEKTQDQQSLSDVEGAYSRAEATRGAGGSSSSPPEKDSGLLDS |

| | |
|---|---|
| | VLDTLLAPSGPGQSQPSPPACEVTSSWCLFGPELPEDPPAAPATQRVLSPLMSRSGC KVGDSSGTAAAHKVLPRGLSPARQLLLPASESPHWSGAPVKPSPQAAAVEVEEEDG SESEESAGPLLKGKPRALGGAAAGGGAAAVPPGAAAGGVALVPKEDSRFSAPRVA LVEQDAPMAPGRSPLATTVMDFIHVPILPLNHALLAARTRQLLEDESYDGGAGAAS AFAPPRSSPCASSTPVAVGDFPDCAYPPDAEPKDDAYPLYSDFQPPALKIKEEEEGA EASARSPRSYLVAGANPAAFPDFPLGPPPPLPPRATPSRPGEAAVTAAPASASVSSAS SSGSTLECILYKAEGAPPQQGPFAPPPCKAPGASGCLLPRDGLPSTSASAAAAGAAP ALYPALGLNGLPQLGYQAAVLKEGLPQVYPPYLNYLRPDSEASQSPQYSFESLPQKI $^{566}$ |
| GR AD | $^{1}$MDSKESLTPGREENPSSVLAQERGDVMDFYKTLRGGATVKVSASSPSLAVASQSD SKQRRLLVDFPKGSVSNAQQPDLSKAVSLSMGLYMGETETKVMGNDLGFPQQGQI SLSSGETDLKLLEESIANLNRSTSVPENPKSSASTAVSAAPTEKEFPKTHSDVSSEQQ HLKGQTGTNGGNVKLYTTDQSTFDILQDLEFSSGSPGKETNESPWRSDLLIDENCLL SPLAGEDDSFLLEGNSNEDCKPLILPDTKPKIKDNGDLVLSSPSNVTLPQVKTEKEDF IELCTPGVIKQEKLGTVYCQASFPGANIIGNKMSAISVHGVSTSGGQMYHYDMNTA SLSQQQDQKPIFNVIPPIPVGSENWNRCQGSGDDNLTSLGTLNFPGRTVFSNGYSSPS MRPDVSSPPSSSSTATTGPPPKL$^{420}$ |
| ERα AD | $^{1}$MTMTLHTKASGMALLHQIQGNELEPLNRPQLKIPLERPLGEVYLDSSKPAVYNYPE GAAYEFNAAAAANAQVYGQTGLPYGPGSEAAAFGSNGLGGFPPLNSVSPSPLMLL HPPPQLSPFLQPHGQQVPYYLENEPSGYTVREAGPPAFYRPNSDNRRQGGRERLAS TNDKGSMAMESAKETRY$^{184}$ |
| Nt | $^{1}$MEVQLGLGRVYPRPPSKTYRGAFQNLFQSVREVIQNPGPRHPEAASAAPPGASLLL LQQQQQQQQQQQQQQQQQQQQQETSPRQQQQQQGEDGSPQAHRRGPTGYLVLD EEQQPSQPQSALECHPERGCVPEPGAAVAASKGLPQQLPAPP$^{151}$ |
| Nt C123 (C129S) | $^{1}$MEVQLGLGRVYPRPPSKTYRGAFQNLFQSVREVIQNPGPRHPEAASAAPPGASLLL LQQQQQQQQQQQQQQQQQQQQQETSPRQQQQQQGEDGSPQAHRRGPTGYLVLD EEQQPSQPQSALECHPERG**S**VPEPGAAVAASKGLPQQLPAPP$^{151}$ |
| Nt L26P | $^{1}$MEVQLGLGRVYPRPPSKTYRGAFQN**P**FQSVREVIQNPGPRHPEAASAAPPGASLLL LQQQQQQQQQQQQQQQQQQQQQETSPRQQQQQQGEDGSPQAHRRGPTGYLVLD EEQQPSQPQSALECHPERGCVPEPGAAVAASKGLPQQLPAPP$^{151}$ |
| Tau-5* | $^{330}$AAGSSGTLELPSTLSLYKSGALDEAAAYQSRDYYNFPLALAGPPPPPPPPHPHARI KLENPLDYGSAWAAAAAQCRYGDLASLHGAGAAGPGSGSPSAAASSSWHTLFTA EEGQLYGPC$^{448}$ |
| Tau-5* C404 (C448 deleted) | $^{330}$AAGSSGTLELPSTLSLYKSGALDEAAAYQSRDYYNFPLALAGPPPPPPPPHPHARI KLENPLDYGSAWAAAAAQCRYGDLASLHGAGAAGPGSGSPSAAASSSWHTLFTA EEGQLYGP*****$^{448}$ |
| Tau-5* C404S | $^{330}$AAGSSGTLELPSTLSLYKSGALDEAAAYQSRDYYNFPLALAGPPPPPPPPHPHARI KLENPLDYGSAWAAAAAQ**S**RYGDLASLHGAGAAGPGSGSPSAAASSSWHTLFTAE EGQLYGPC$^{448}$ |

| | |
|---|---|
| noAro (all aromatic residues are substituted with A) | $^{330}$AAGSSGTLELPSTLSL**A**KSGALDEAAA**A**QSRD**AA**N**A**PLALAGPPPPPPPP**A**P**A**ARI<br>KLENPLD**A**GSA**A**AAAAAQCR**A**GDLASL**A**GAGAAGPGSGSPSAAASSS**AA**TL**A**TAE<br>EGQL**A**GPC$^{448}$ |
| h2SA (Y393S+W397A+ Y406S+H413A) | $^{330}$AAGSSGTLELPSTLSLYKSGALDEAAAYQSRDYYNFPLALAGPPPPPPPPHPHARI<br>KLENPLD**S**GSA**A**AAAAAQCR**S**GDLASL**A**GAGAAGPGSGSPSAAASSSWHTLFTAE<br>EGQLYGPC$^{448}$ |
| h3A (W433A+H434A+ F437A) | $^{330}$AAGSSGTLELPSTLSLYKSGALDEAAAYQSRDYYNFPLALAGPPPPPPPPHPHARI<br>KLENPLDYGSAWAAAAAQCRYGDLASLHGAGAAGPGSGSPSAAASSS**AA**TL**A**TAE<br>EGQLYGPC$^{448}$ |
| C404Y | $^{330}$AAGSSGTLELPSTLSLYKSGALDEAAAYQSRDYYNFPLALAGPPPPPPPPHPHARI<br>KLENPLDYGSAWAAAAAQ**Y**RYGDLASLHGAGAAGPGSGSPSAAASSSWHTLFTA<br>EEGQLYGPC$^{448}$ |
| AAA (G394A+S395A+ G407A) | $^{330}$AAGSSGTLELPSTLSLYKSGALDEAAAYQSRDYYNFPLALAGPPPPPPPPHPHARI<br>KLENPLDY**AA**AWAAAAAQCRYADL**A**SLHGAGAAGPGSGSPSAAASSSWHTLFTA<br>EEGQLYGPC$^{448}$ |
| A398P | $^{330}$AAGSSGTLELPSTLSLYKSGALDEAAAYQSRDYYNFPLALAGPPPPPPPPHPHARI<br>KLENPLDYGSAW**P**AAAAQCRYGDLASLHGAGAAGPGSGSPSAAASSSWHTLFTAE<br>EGQLYGPC$^{448}$ |
| L436P | $^{330}$AAGSSGTLELPSTLSLYKSGALDEAAAYQSRDYYNFPLALAGPPPPPPPPHPHARI<br>KLENPLDYGSAWAAAAAQCRYGDLASLHGAGAAGPGSGSPSAAASSSWHT**P**FTA<br>EEGQLYGPC$^{448}$ |
| T435P | $^{330}$AAGSSGTLELPSTLSLYKSGALDEAAAYQSRDYYNFPLALAGPPPPPPPPHPHARI<br>KLENPLDYGSAWAAAAAQCRYGDLASLHGAGAAGPGSGSPSAAASSSWH**P**LFTA<br>EEGQLYGPC$^{448}$ |
| A398P+L436P | $^{330}$AAGSSGTLELPSTLSLYKSGALDEAAAYQSRDYYNFPLALAGPPPPPPPPHPHARI<br>KLENPLDYGSAW**P**AAAAQCRYGDLASLHGAGAAGPGSGSPSAAASSSWHT**P**FTAE<br>EGQLYGPC$^{448}$ |
| A398P+T435P | $^{330}$AAGSSGTLELPSTLSLYKSGALDEAAAYQSRDYYNFPLALAGPPPPPPPPHPHARI<br>KLENPLDYGSAW**P**AAAAQCRYGDLASLHGAGAAGPGSGSPSAAASSSWH**P**LFTAE<br>EGQLYGPC$^{448}$ |
| G407A | $^{330}$AAGSSGTLELPSTLSLYKSGALDEAAAYQSRDYYNFPLALAGPPPPPPPPHPHARI<br>KLENPLDYGSAWAAAAAQCRY**A**DLASLHGAGAAGPGSGSPSAAASSSWHTLFTA<br>EEGQLYGPC$^{448}$ |
| Ct | $^{441}$EGQLYGPCGGGGGGGGGGGGGGGGEAGAVAPYGYTRPPQGLAG<br>QESDFTAPDVWYPGGMVSRVPYPSPTCVKSEMGPWMDSYSGPYGDMRLETARDH<br>VLPIDYYFPPQKT$^{558}$ |

| | |
|---|---|
| Ct 4G | 441 EGQLYGPCGG********************GGEAGAVAPYGYTRPPQGLAGQESDFTA PDVWYPGGMVSRVPYPSPTCVKSEMGPWMDSYSGPYGDMRLETARDHVLPIDYY FPPQKT 558 |
| Ct C518 (C448 is deleted) | 441 EGQLYGP*GG********************GGEAGAVAPYGYTRPPQGLAGQESDFTA PDVWYPGGMVSRVPYPSPTCVKSEMGPWMDSYSGPYGDMRLETARDHVLPIDYY FPPQKT 558 |
| allTau5 | 330 AAGSSGTLELPSTLSLYKSGALDEAAAYQSRDYYNFPLALAGPPPPPPPPHPHARI KLENPLDYGSAWAAAAAQCRYGDLASLHGAGAAGPGSGSPSAAASSSWHTLFTA EEGQLYGPCGG********************GGEAGAVAPYGYTRPPQGLAGQESDFTAP DVWYPGGMVSRVPYPSPTCVKSEMGPWMDSYSGPYGDMRLETARDHVLPIDYYF PPQKT 558 |
| allTau-5 CtoS | 330 AAGSSGTLELPSTLSLYKSGALDEAAAYQSRDYYNFPLALAGPPPPPPPPHPHARI KLENPLDYGSAWAAAAAQSRYGDLASLHGAGAAGPGSGSPSAAASSSWHTLFTAE EGQLYGPSGG********************GGEAGAVAPYGYTRPPQGLAGQESDFTAPD VWYPGGMVSRVPYPSPTSVKSEMGPWMDSYSGPYGDMRLETARDHVLPIDYYFPP QKT 558 |
| allTau-5 CtoS PP | 330 AAGSSGTLELPSTLSLYKSGALDEAAAYQSRDYYNFPLALAGPPPPPPPPHPHARI KLENPLDYGSAWPAAAAQSRYGDLASLHGAGAAGPGSGSPSAAASSSWHTPFTAE EGQLYGPSGG********************GGEAGAVAPYGYTRPPQGLAGQESDFTAPD VWYPGGMVSRVPYPSPTSVKSEMGPWMDSYSGPYGDMRLETARDHVLPIDYYFPP QKT 558 |
| allTau-5 C404 | 330 AAGSSGTLELPSTLSLYKSGALDEAAAYQSRDYYNFPLALAGPPPPPPPPHPHARI KLENPLDYGSAWAAAAAQCRYGDLASLHGAGAAGPGSGSPSAAASSSWHTLFTA EEGQLYGPSGG********************GGEAGAVAPYGYTRPPQGLAGQESDFTAP DVWYPGGMVSRVPYPSPTSVKSEMGPWMDSYSGPYGDMRLETARDHVLPIDYYF PPQKT 558 |
| allTau-5 C448 | 330 AAGSSGTLELPSTLSLYKSGALDEAAAYQSRDYYNFPLALAGPPPPPPPPHPHARI KLENPLDYGSAWAAAAAQSRYGDLASLHGAGAAGPGSGSPSAAASSSWHTLFTAE EGQLYGPCGG********************GGEAGAVAPYGYTRPPQGLAGQESDFTAP DVWYPGGMVSRVPYPSPTSVKSEMGPWMDSYSGPYGDMRLETARDHVLPIDYYF PPQKT 558 |
| allTau-5 C518 | 330 AAGSSGTLELPSTLSLYKSGALDEAAAYQSRDYYNFPLALAGPPPPPPPPHPHARI KLENPLDYGSAWAAAAAQSRYGDLASLHGAGAAGPGSGSPSAAASSSWHTLFTAE EGQLYGPSGG********************GGEAGAVAPYGYTRPPQGLAGQESDFTAPD VWYPGGMVSRVPYPSPTCVKSEMGPWMDSYSGPYGDMRLETARDHVLPIDYYFP PQKT 558 |

* amino acid deletion.

**Table S2.** Primers used for the amplification of constructs in the LacI tethering assay.

| | |
|---|---|
| CACCGGGTTCTGCGGGTTCTGCCGCAGGTggatccGCGATaATGACCATGACCCTCCACACCAAAGCATCTGGGATGGCC | pJM118-CFP-LacI-ER_AD_Human-FW |
| gcaagcttgtcgacggcgctcgaattcGGGCCCTCTAGACtcaGTAGCGAGTCTCCTTGGCAGAT | pJM118-CFP-LacI-ER_AD_Human-RV |
| CACCGGGTTCTGCGGGTTCTGCCGCAGGTggatccGCGATaATGGAGACCAAAGGCTACCACAGT | pJM118-CFP-LacI-NR3C2(MR)_AD_Human-FW |
| gcaagcttgtcgacggcgctcgaattcGGGCCCTCTAGACTCATATTTTTGAAGGTCTTGAAGATCCAGTAGAAACACTT | pJM118-CFP-LacI-NR3C2(MR)_AD_Human-RV |
| CACCGGGTTCTGCGGGTTCTGCCGCAGGTggatccGCGATaATGGACTCCAAAGAATCATTAACTCCTGGTAGAGAAGAAAACCCCAG | pJM118-CFP-LacI-NR3C1(GR)_AD_Human-FW |
| gcaagcttgtcgacggcgctcgaattcGGGCCCTCTAGACtcaGAGTTTGGGAGGTGGTCCTGT | pJM118-CFP-LacI-NR3C1(GR)_AD_Human-RV |
| CACCGGGTTCTGCGGGTTCTGCCGCAGGTggatccGCGATaATGACTGAGCTGAAGGCAAAGG | pJM118-CFP-LacI-PGR(PR)_AD_Human-FW |
| gcaagcttgtcgacggcgctcgaattcGGGCCCTCTAGACtcaAATCTTCTGAGGTAATGACTCGAAGCTG | pJM118-CFP-LacI-PGR(PR)_AD_Human-RV |
| CACCGGGTTCTGCGGGTTCTGCCGCAGGTggatccGCGATaATGGAAGTGCAGTTAGGGCT | pJM118-CFP-LacI-AR_AD_Human-FW |
| gcaagcttgtcgacggcgctcgaattcGGGCCCTCTAGACtcaGGTCTTCTGGGGTGGAAAGTAATAGTCA | pJM118-CFP-LacI-AR_AD_Human-RV |
| CACCGGGTTCTGCGGGTTCTGCCGCAGGTggatccGCGATaGCGTCCAATGACTACACTCAGCAG | pJM118_hFUS-LCD-FW |
| gcaagcttgtcgacggcgctcgaattcGGGCCCTCTAGACTCAGATGGTGTT | pJM118_hFUS-LCD-RV |
| CACCGGGTTCTGCGGGTTCTGCCGCAGGTggatccGCGATaATGACTGAGCTGAAGGCAAAGG | pJM118-CFP-LacI-PGR(PR)_AD_Human-WF |
| gcaagcttgtcgacggcgctcgaattcGGGCCCTCTAGACtcaAATCTTCTGAGGTAATGACTCGAAGCTG | pJM118-CFP-LacI-PGR(PR)_AD_Human-RV |

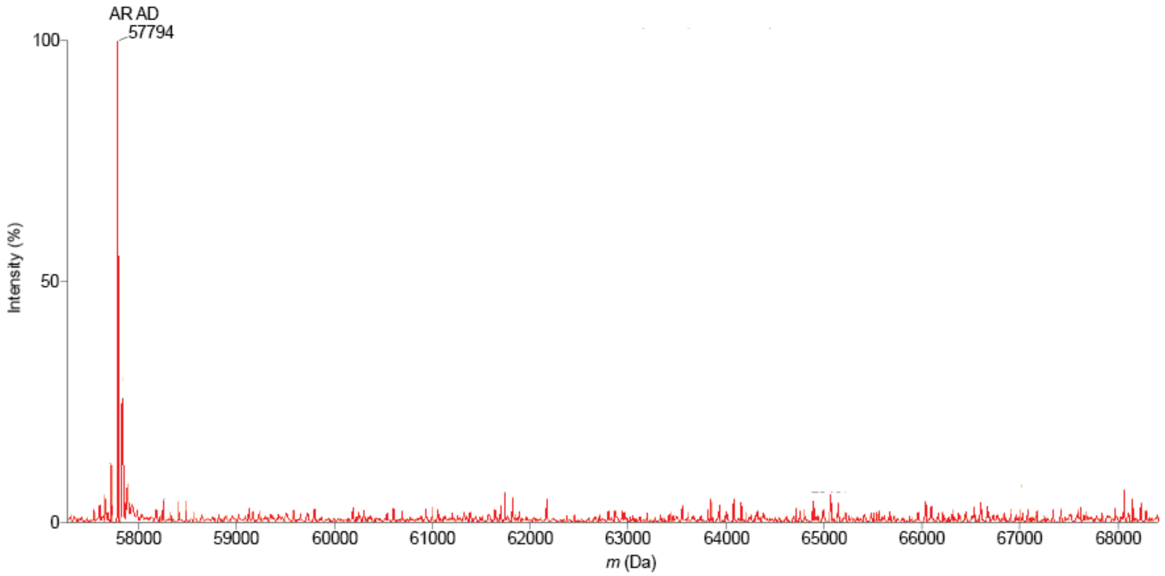


**Fig. S1** Deconvoluted mass spectra of AR AD incubated with EPI-001 for 48 hours under conditions consistent with those used for NMR studies of EPI-001 binding: pH 7.4, 5 °C. It was not detected peaks corresponding to +358.43n Da, where n represents the number of cysteines reacted with EPI-001, AR AD contains 11 cysteines.

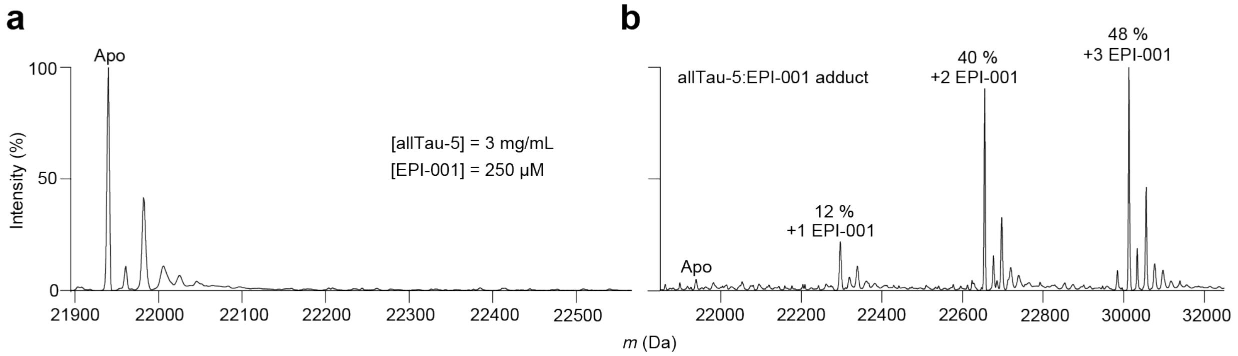


**Fig. S2 (a)** Deconvoluted mass spectra of allTau-5 incubated with EPI-001 for 30 minutes at conditions that were used to study the condensates by fluorescence microscopy. No protein adduct was observed, indicating that the observed effects report exclusively on the reversible binding. **(b)** Deconvoluted mass spectra of allTau-5:EPI-001 adduct, revealing covalent interaction of EPI-001 with all three cysteines in allTau5.

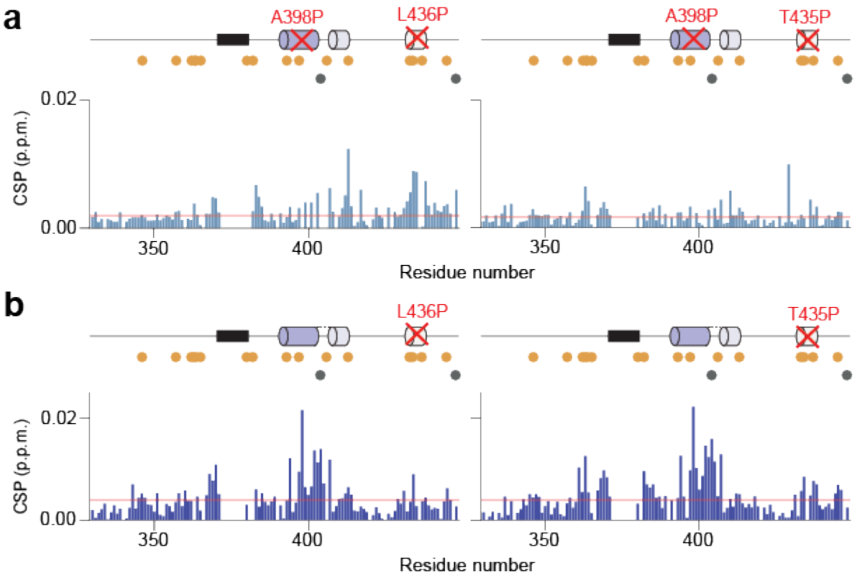


**Fig. S3.** (a) CSPs detected in 2D $^{1}$H-$^{15}$N NMR correlation spectra of Tau-5* mutants A398P+L436P and A398P+T435P by increasing the protein concentration from 200 µM to 400 µM. (b) CSPs observed in the NMR spectra of Tau-5* mutants L436P and T435P by increasing the protein concentration from 25 µM to 400 µM. Orange and grey dots indicate the positions of aromatic and cysteine residues, respectively. The red line represents the significant threshold calculated as the average plus five standard deviations of the first quartile of CSPs.

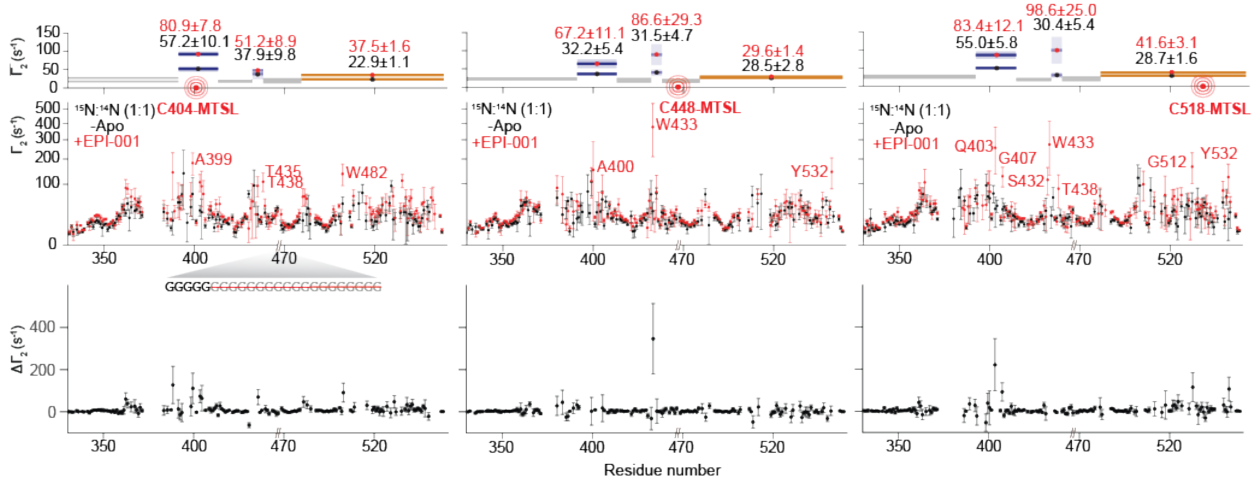


**Fig. S4** (*Top*) Intermolecular interactions between allTau5 molecules monitored via PRE NMR experiments in the absence (black) and presence of 1 molar equivalent of the ligand (red). (*Bottom*) Difference in PREs observed in the presence versus absence of EPI-001. Error bars indicate the standard error from the exponential fit. The average $\Gamma_2^{HN}$ for selected regions are shown in upper plots. Error bars represent the standard deviations.

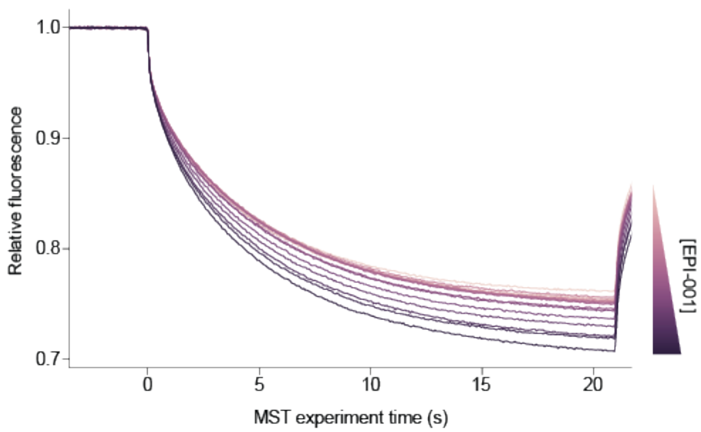


**Fig. S5** MST traces of 80 nM 2Tau-5* in the presence of increased concentrations of EPI-001 (from 1.6 to 500 μM).
